# ribofootPrinter: A precision python toolbox for analysis of ribosome profiling data

**DOI:** 10.1101/2021.07.04.451082

**Authors:** Kyra Kerkhofs, Nicholas R. Guydosh

**Affiliations:** Laboratory of Biochemistry and Genetics, National Institute of Diabetes and Digestive and Kidney Diseases, National Institutes of Health, Bethesda, MD, 20892

**Keywords:** Ribosome profiling, reading frame, metagene, pausing, uORF, iORF, dORF, Python, MANE, multimapping

## Abstract

Ribosome profiling is a valuable methodology for measuring changes in a cell’s translational program. The technique can report how efficiently mRNA coding sequences are translated and pinpoint positions along mRNAs where ribosomes slow down or arrest. It can also reveal when translation takes place outside coding regions, often with important regulatory consequences. While many useful software tools have emerged to facilitate analysis of these data, packages can become complex and challenging to adapt to specialized needs. We therefore introduce ribofootPrinter, a suite of Python tools designed to offer an accessible and modifiable set of code for analysis of data from ribosome profiling and related types of small RNA sequencing experiments. Read alignments are made to a simplified transcriptome to keep the code intuitive. Multiple normalization options help facilitate interpretation of data, particularly outside coding regions. We also demonstrate how the length of reads that map to the transcriptome affects the frequency of matches to multiple sites and we provide multimapper identifier files to highlight these regions. Overall, this tool has the capability to carry out sophisticated analyses while maintaining enough simplicity to make it readily understandable and adaptable.

## Introduction

Ribosome profiling (also referred to as Ribo-seq) has become an established approach for revealing which genes are translated and how ribosomes are distributed along transcripts in cells (Ingolia et al. 2009; McGlincy and Ingolia 2017). Ribosome profiling data are generated by deep sequencing cDNA libraries that are generated from ribosome-protected mRNA footprints that typically measure ∼28-30 nt in length (Figure 1A). However, in some experimental conditions, footprints can measure ∼16 nt, ∼21 nt, or >29 nt (Lareau et al. 2014; Schmidt et al. 2016; Guydosh and Green 2017; Wu et al. 2019; Ganesan et al. 2022) and reveal additional information. To analyze ribosome profiling datasets, reads are typically aligned to the transcriptome and software packages are used to reveal a rich and precise view of a cell’s translational program. Analysis can show the distribution and loading (translational efficiency) of ribosomes across messenger RNAs (mRNAs). It can also offer insight on where actively-translated open reading frames (ORFs) exist, including within non-canonical regions, such as 5’ and 3’ untranslated regions (UTRs) and out-of-frame sequences within coding sequences (CDSs) (Mudge et al. 2022). These small ORFs are commonly referred to as upstream ORFs (uORFs), downstream ORFs (dORFs) or internal ORFs (iORFs), respectively. Related approaches (Dunn and Weissman 2016; Liu et al. 2020; Tjeldnes et al. 2021) for footprinting disomes (disome-seq) or 40S ribosomal subunits (TCP-seq) (Archer et al. 2016; Bohlen et al. 2020; Meydan and Guydosh 2020; Young et al. 2021) have enhanced the method to address questions related to ribosome collisions and translation initiation, respectively.

**Figure 1.**
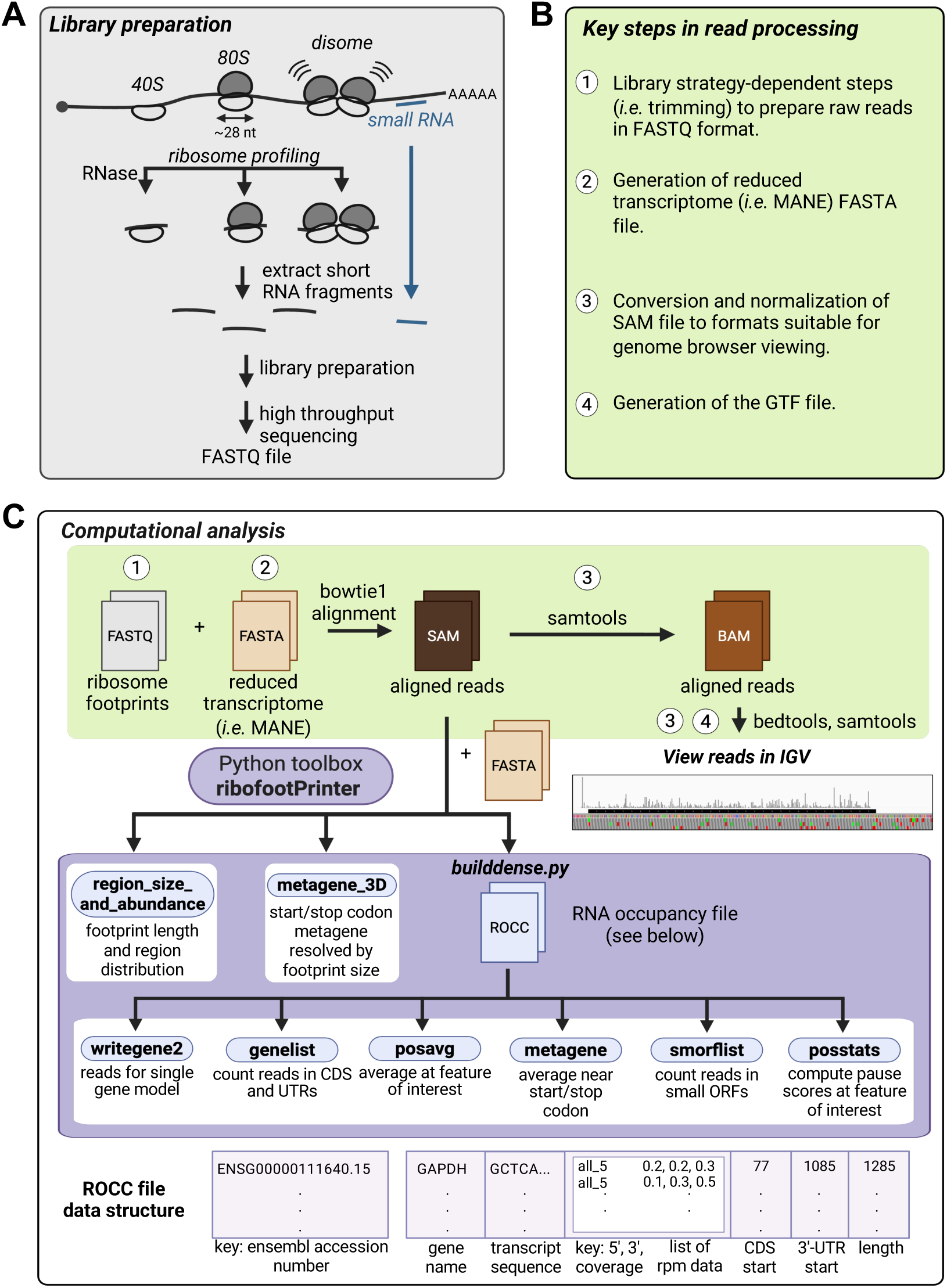
ribofootPrinter is a comprehensive Python toolbox for analysis of short RNA datasets. **(A)** Schematic overview of short RNA datasets compatible with ribofootPrinter. Short RNA libraries are sequenced to generate high-throughput datasets in FASTQ format. **(B)** List of major steps in read processing prior to running key scripts in ribofootPrinter. **(C)** Steps to analyze data with ribofootPrinter. Pre-processed reads in FASTQ format are first aligned to the reduced transcriptome (FASTA) with bowtie1 (*top*). The resultant SAM file is processed into an RNA occupancy file (ROCC) using the *builddense* script. The ROCC file data structure contains ribosome occupancy data (5’- or 3’-assigned, or coverage) and sequence metadata (*bottom*). Data can then be analyzed with six downstream analysis tools: *writegene2, genelist, posavg, metagene, smorflist,* and *posstats*. Both *region_size_and_abundance* and *metagene_3D* are directly compatible with SAM files and do not require a ROCC file. Steps to view data in the IGV browser include conversion of SAM files into IGV-compatible BAM and BEDGRAPH files (*top right*).

Many capable software tools have emerged to analyze ribosome profiling data, in part because no single approach fits every need (Limbu et al. 2024). However, one of the challenges faced by all approaches is how to handle the complex nature of mRNA splicing patterns in the cell. In humans, most transcripts are spliced into multiple isoforms, and this creates the problem of how to represent a particular gene’s level of ribosome occupancy, particularly when different isoforms (gene models) are translated with varied efficiency. Several solutions have emerged, including use of masking functions to exclude transcript regions where splicing patterns are variable (Dunn and Weissman 2016). Other approaches simplify the problem by mapping to the longest isoform or isoform with the most exons (Ishimura et al. 2014; Sendoel et al. 2017; Ahmed et al. 2019). Another challenge posed by splicing is the computational burden of converting between genomic and transcriptomic coordinate systems (Tjeldnes et al. 2021). This complexity also tends to make the code difficult to understand, raising barriers to customization of the code for new users. Recently, the Matched Annotation from the NCBI and EMBL-EBI (MANE) project offered a curated transcriptome for *Homo sapiens* with a single recognized (most likely to be used) isoform per gene (Morales et al. 2022). This unique resource offers a simple way to solve the problems associated with splicing for users who are not particularly concerned with unique isoforms.

Another challenge at the alignment step is deciding how to assign reads that can map to multiple locations in the transcriptome. While some allowance for multiple mapping locations is typical in bioinformatic pipelines, there is a limit to it and this becomes particularly important when reads are short. As a result, it is important to use caution when interpreting data that map to transcriptome regions with high degeneracy.

Once ribosome profiling data are aligned, the next challenge is undertaking more specialized analysis that includes assessment of where reads are enriched. A common approach is to create a “meta” plot by averaging ribosome profiling data over particular sequence motifs of interest (*i.e.* metagene or metacodon analysis). The simplest form of this analysis involves computing the arithmetic mean (or median) for read counts at every nucleotide position surrounding the feature of interest. However, the downside of this approach is that every feature (*i.e.* gene or codon) is not weighted equally in the average; highly expressed features tend to dominate. The need to consider alternatives to normalization is particularly critical when considering translation outside conventional coding sequences (Chen et al. 2020; Karasik et al. 2021; Mudge et al. 2022). In such cases, the most appropriate method of normalization may not be immediately obvious, and it is therefore advantageous to consider multiple forms of normalization. Normalization considerations can also impact related analyses of read count distributions within particular regions of the transcriptome.

A final challenge of many computational tools is the loss of support over time and their eventual deprecation as changes are inevitably made to supporting packages and the programming languages themselves. This problem is particularly acute for packages that are complex and rely on extensive outside software to function. Therefore, simple and readily modifiable tools are advantageous for promoting longevity and continuity.

We therefore introduce ribofootPrinter, a precision Python toolbox for analysis of ribosome profiling data that can be adapted to analyze data from any organism. The package simplifies the splicing problem by relying on a spliced transcriptome (for example, provided by the MANE project) and offers several functions that can be readily modified. Importantly, the software also offers the option of multiple normalization approaches for meta-analysis and comparison of read abundance across transcriptome regions. The simplicity of the approach makes this package appealing for both introductory programmers as well as users interested in a straightforward platform. ribofootPrinter is particularly useful for understanding several topics of growing importance, such as finding translationally active small ORFs, assessing ribosome stalling at sequences of interest, deconvolving variation in footprint lengths within particular regions, and evaluating translation outside of a transcript’s annotated coding sequence (CDS), which we also refer to as the main open reading frame (ORF). The tool is compatible with 80S conventional ribosome footprinting, as well as 40S and disome footprinting, and can be adapted to other forms of small RNA sequencing.

### Implementation

#### Pipeline overview

We provide a complete and comprehensive analysis pipeline (Figure 1) for datasets obtained from ribosome profiling and other methods that use Illumina sequencing, such as sequencing of miRNA, piRNA, or CLIP-derived RNA. The first step is to pre-process the high-throughput sequencing FASTQ data file, which contains the raw sequencing reads (Step 1 in Figure 1B, C). Often this includes read trimming, deduplication, and removal of contaminating (for example, rRNA) reads, as described elsewhere (McGlincy and Ingolia 2017).

Next, to determine where reads map in the transcriptome, the processed FASTQ file must be aligned to the reduced transcriptome FASTA file (Figure 1C). We include instructions for using bowtie1 (Langmead et al. 2009), but other aligners can be used. Modification of the MANE version 1.4 (human) transcriptome to be compatible with ribofootPrinter is described in detail in the Methods (Step 2 in Figures 1B, C, and Supplementary Figure 1A). This transcriptome contains 19,404 transcripts but we excluded MANE Select non-coding RNAs to focus on protein coding transcripts (Figure 2A). In addition, we also excluded the MANE Plus Clinical set to avoid duplicate gene names and sequences. This results in a transcriptome with reduced multimapping (*i.e.* many sequences that a single read could be aligned to). For example, the *GNAS* gene (ENSG00000087460) has two isoforms that are annotated as MANE Plus Clinical and are therefore eliminated in this process (Figure 2B). These removals leave a total of 19,288 transcripts for alignment (listed in shortnames.fasta, see Figure Supplementary Figure 1A and Methods).

**Figure 2.**
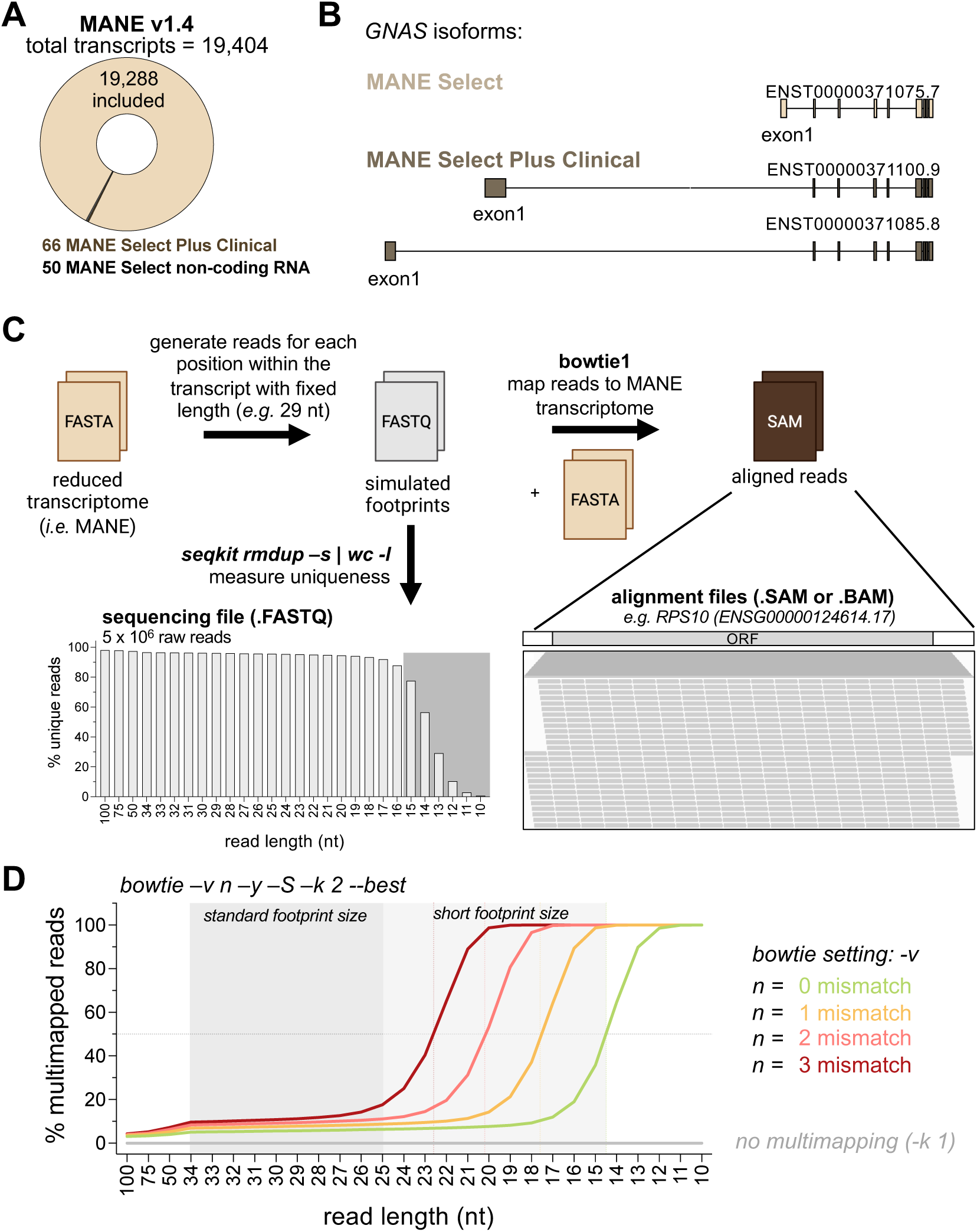
Multimapping depends on read length and number of mismatches. **(A)** Overview of transcripts included in the MANE transcriptome in this study. Clinical and non-coding RNA sequences were excluded to reduce duplicate sequences. **(B)** Schematic representation of multiple isoforms of the *GNAS* transcript. The MANE select transcript isoform is shown in light brown and additional clinical select isoforms are shown in dark brown. **(C)** The reduced MANE v1.4 transcriptome was used to generate reads for each possible position within the transcript for a given length. These transcriptome-derived FASTQ files were aligned back to the exact same reduced transcriptome to obtain aligned SAM files. An example of the IGV browser is shown. Note the complete coverage of the entire RPS10 transcript. The seqkit package was used to determine uniqueness of reads as a function of read length (bar graph). **(D)** Identification of multimapping events following alignment of transcriptome-derived FASTQ files against the reduced transcriptome using bowtie. Different *-v* settings allow for different number of mismatches during alignment. As expected, the FASTQ files containing shorter read lengths are more prone to multimapping while larger read lengths result in more uniquely mapped reads. Increasing the number of mismatches by changing the setting (*-v*) results in increasing multimapped reads.

The SAM file that is generated by the alignment can then be converted into a BAM file and then a BEDGRAPH or BIGWIG file that is normalized by sequencing depth (*i.e.* units of RPM) for visualization in a genome browser, such as IGV (Thorvaldsdottir et al. 2013) (Step 3 in Figures 1B,C, and Supplementary Figure 1B). The BAM file can also be directly viewed to observe individual reads (Supplementary Figure 1B). Unlike traditional approaches where RNA reads are aligned to the genome, this transcriptome alignment produces tracks that lack introns. To generate transcript-based feature annotations, such as alternate names and ORF boundaries, a GTF file (see Methods for file preparation guidance) is loaded by the genome browser for data visualization (Step 4 in Figures 1B,C, and Supplementary Figure 1B). Additionally, the SAM file can be converted into a specialized RNA occupancy (“ROCC”) file that is used for downstream, in-depth analysis using ribofootPrinter (Figure 1C). The code is written in Python 3 and requires the BioPython package (Cock et al. 2009) for sequence manipulations. All functionalities typically run within a few minutes on modern personal computers.

While the utilization of a reduced transcriptome represents an obvious compromise, it facilitates downstream analysis by simplifying the code and eliminates artifacts associated with using the longest (but not most common) transcript isoform or eliminating common genes due to ambiguous splicing. This curation process also avoids non-ribosome derived peaks sometimes found on non-coding RNAs. In addition, it allows easy visualization of translation events in genome browsers, such as IGV, without interference from introns.

#### Reduced transcriptome alignment considerations

To test basic properties of read alignment to the MANE 1.4 transcriptome, particularly the tendency to map to multiple locations, we generated a FASTQ file containing every possible read that could be derived from the transcriptome sequences (Figure 2C). Since ribosomes can protect a range of mRNA lengths, depending on preparation conditions, we generated files for footprint sizes ranging from 10-100 nt. These lengths also span the typical lengths of other small RNAs of interest, such as miRNAs and piRNAs. We found that 2% of the 100-mer reads were not unique (Figure 2C). This result could potentially be attributed to repetitive sequences, conservation of sequences between genes, different isoforms of the same gene, or overlapping transcripts (Veeramachaneni et al. 2004). As expected, the number of unique reads declined as read length (and therefore information content) was reduced and fell off steeply below about 15 nucleotides) (Figure 2C, fastq file bar graph gray background).

Next, we aligned these transcriptome-derived FASTQ files back to the transcriptome using bowtie (Figure 2C) to explore how settings in the alignment affect this process. We confirmed that 100% of the reads mapped following bowtie mapping using different settings. Consistent with our finding that shorter (particularly <15 nt) reads contain a large proportion of non-unique sequences (see Figure 2C), we found that the fraction of reads mapping to multiple locations increased for shorter read lengths (Figure 2D). To maximize the number of reads that mapped uniquely, we specified 0 mismatches in the alignment (*-v 0*) to ensure perfect alignment of the read to the transcriptome. With this setting, we found 65% of the 15-mer reads mapped uniquely (35% mapped to multiple locations) (Figure 2D) while, in contrast, 90% of 20-mer reads mapped uniquely. We noted this trend changed as the number of mismatches was relaxed, going from 8% with 0 mismatches, 14% with 1 mismatch, 53% with 2 mismatches and 99% with 3 mismatches for the 20-mer reads (Figure 2D). Another consideration is that since the transcriptome only contains positive-sense annotated transcripts, running alignments that disallow mapping to the reverse complement strand *(--norc*; no reverse complement) enhanced performance (6 vs 8% multimapping with 0 mismatches for 20-mer reads) (data not shown). Whether limiting antisense mapping is advantageous may depend on the particular conditions of the experiment. For example, a tendency for some bidirectionality in many promoters may contribute to antisense reads in real-world datasets (Nemsick and Hansen 2024).

To help characterize the inherent repetitiveness of the transcriptome that underlies the tendency of reads to map to multiple locations, we generated multimapper identifier files (*mm_id.bedgraph)* for a range of read lengths to highlight these regions in a genome browser (Supplementary Figure 2). The metric encoded in these files measures the tendency for a read to map to multiple locations with higher values indicating more multimapping (see Methods).

To demonstrate the utility of the multimapper identifier files, we examined the *ACTB* transcript (encoding β-actin). We noted several regions that included clusters of multiply mapping reads (Figure 3A, highlighted in grey). We identified two transcripts, *POTEI* and *POTEJ*, that contain a high level of sequence conservation with *ACTB* and are responsible for the non-unique mapping (Figure 3B) (Madeira et al. 2024). Consistent with the analysis above, the tracks made for longer read lengths contain lower values compared to those made for shorter reads. When we examined our Ribo-Seq alignment where limited multimapping was tolerated (multi-mapping reads assigned to one location at random, bowtie setting *-k 1*), we noted generally fewer mapped ribosome footprints within segments with higher multimapper scores compared to the surrounding regions (Figure 3A, top row). We further noted that the *POTEI* and *POTEJ* transcripts generally had no footprints mapping to them except for the multimapped region (Figure 3C). We therefore conclude that reads from these regions likely derive from *ACTB* and are evenly divided between three transcripts, resulting in diminished mapping to *ACTB* and unexpected mapping to *POTEI* and *POTEJ*. As a result, performing ribofootPrinter’s pause score analysis (see later) on this region for proline, which is known to induce ribosome pausing (Schuller et al. 2017), would tend to fail to identify the pause (Figure 3A, zoom). While the examples of non-unique sequences within *ACTB* arise from small regions of homology, we note other examples where entire exons are conserved between many genes in the MANE transcriptome. An example is the protocadherin protein family that is annotated with the *PCDHG* prefix. The multi-mapping problem has been long established in the bioinformatics field and best approach for handling it (allowing 0, 1, or many locations to be mapped, or adding additional processing) will vary according to the needs of particular applications (Wang et al. 2016; Halpin et al. 2020). Our analysis here is aimed at making these limitations clear as a helpful resource for users.

**Figure 3.**
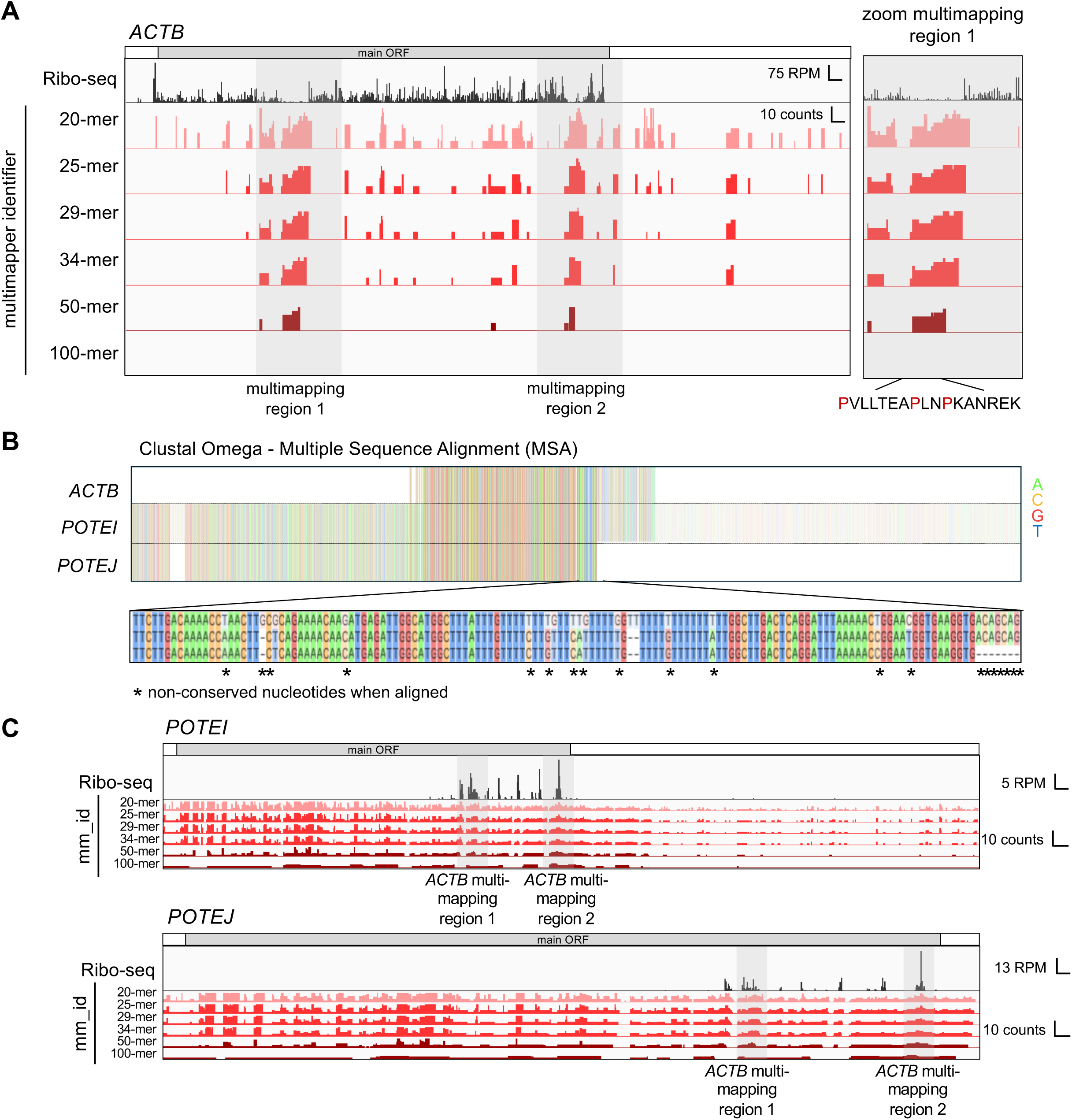
Multimapper identifier files highlight multimapping events on *ACTB*. **(A)** IGV browser view of the *ACTB* transcript (encoding β-actin) with Ribo-seq data (black) and multimapper identifier files (mm_id) loaded in different shades of red. Multimapper identifier files containing larger sequence lengths are less likely to multimap (show in darker shades of red) while shorter sequences lengths are more likely to multimap (shown in lighter shades of red). Regions which are identified as likely to contain non-unique (*i.e.* multimapping) sequences are highlighted with a grey box. The multimapped regions correlate with a decrease in footprints in the Ribo-seq dataset caused by default bowtie settings for alignment which only report one single best alignment in the case of multimapped reads. The zoomed box of multimapped region 1 highlights the presence of prolines which are ribosome-stalling inducing sequences. This could lead to an underestimation of stalling events. Full range of y-axis is 75 RPM for Ribo-seq and 10 counts for multimapper identifiers. **(B)** Clustal Omega alignment of transcripts *ACTB, POTEI* and *POTEJ* reveal strong sequence similarities, leading to multimapping. However, the sequences between *ACTB* and *POTEI* and *POTEJ* are not 100% identical, consistent with the absence of multimapped regions in the 100-mer multimapper identifier file in **A**. **(C)** IGV browser view of *POTEI* and *POTEJ* transcripts with Ribo-seq (black) and mm_id data shown in different shades of red. The multimapping regions identified in *ACTB* in **A** are highlighted in grey boxes and reveal the footprints that are likely derived from *ACTB*. Outside the multimapped regions, very few footprints are observed within the main ORF, indicating that these transcripts are not translated. Full range of y-axis is 5 RPM (top) or 13 RPM (bottom) for Ribo-seq and 10 counts for multimapper identifiers.

#### Distribution of footprint lengths

Following alignment of the FASTQ file containing ribosome footprints to the reduced MANE transcriptome (see Figure 1C), the resulting SAM files containing mapped reads can be used to determine the distribution of read lengths (Figure 4A). This analysis can be helpful in determining a dataset’s quality, for example revealing cases where the RNase treatment was insufficient to fully trim footprints to 28-30 nt. In addition, this function can be used to reveal different species in an experiment, for example short-length (*i.e.* 16 or 21 nt) ribosome footprints, disome or trisome footprints, and 40S footprints are distributed differently by length than traditional 28-30 nt footprints. The package also calculates the abundance of footprints according to the sequence region to which they map (*i.e.* start and stop codon, main ORF, and UTRs). This is done by defining an adjustable window around the start and stop codon and otherwise assigning reads to the main ORF or a UTR region (Supplementary Figure 3, top). The distribution of read lengths can then be computed from each of these region-based assignments. Plotting the data in this way (Figure 4B) reveals that ribosome-protected footprints at the stop codon are consistently 1 nt longer compared to other regions, as reported previously (Ingolia et al. 2011). While the region-specific read length distributions can be plotted as a standard probability density (as shown in Figure 4B), other normalization options are available. The most straightforward option is to report raw reads or reads normalized by library depth (RPM units) to facilitate comparison between samples. This approach generates an abundance distribution (Figure 4C, left) that shows most reads within the coding region (main ORF). An alternative way to represent abundance is to compute the fraction of the read density per unit length. In this way, regions that are short but densely populated with reads, receive higher representation (Figure 4C, right). This approach shows start and stop regions to be highly represented due to a peak on those codons, while the UTRs are less represented due to the low mapping of reads per length in these regions. If desired, the length distribution can be combined with abundance normalization to simultaneously visualize both outputs in a single plot (Supplementary Figure 3 bottom). In addition, a subset or single transcript can be studied by providing a list of Ensembl gene accession numbers (geneid) of interest. For example, we determined the read length distribution and region abundances of the uORF-containing *EIF4G2* gene (Figure 4D, accession number ENSG00000110321.19). The abundance plots (Figure 4D, pie charts) show proportionately more reads in the 5’-UTR for this gene compared to the total transcriptome (Figure 4C). This is consistent with the IGV track of *EIF4G2* where footprints within the uORF region (which is annotated as 5’-UTR) (Figure 4E). The histogram of read lengths is similar, as expected (Figure 4D, right).

**Figure 4.**
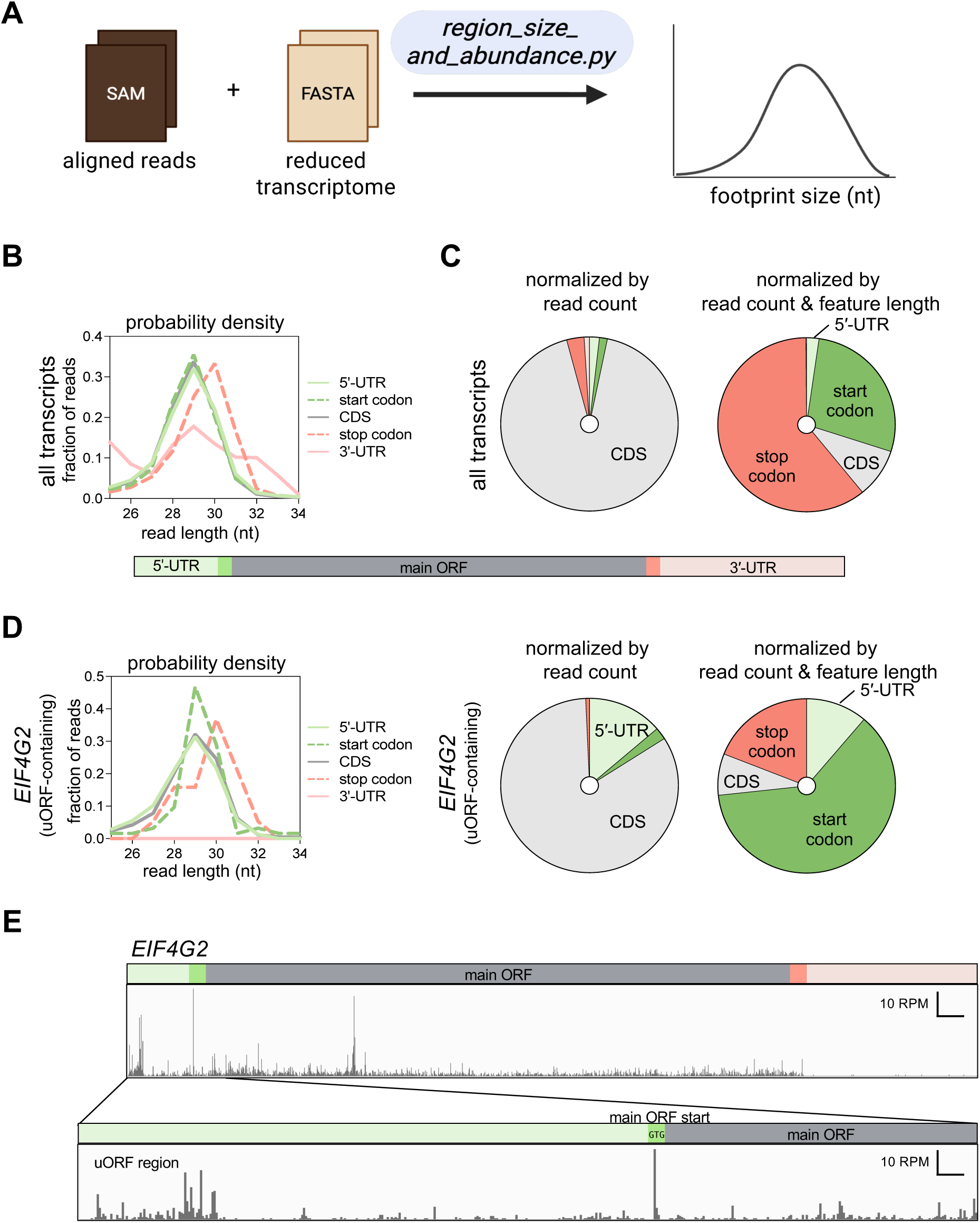
ribofootPrinter analyzes read length distributions within UTRs, main ORF, start codon or stop codon. **(A)** Schematic representation of the *region_size_and_abundance* script. This script uses a SAM file as input. **(B)** The distribution of footprints within UTRs, main ORF, start codon or stop codons for all transcripts normalized by probability density (all sum to 1). This allows direct comparisons between different features, revealing longer stop codon reads in our dataset. **(C)** Abundance of reads across the transcriptome. The pie chart on the left gives the percentage of total reads that map to each region. The pie chart on the right gives the percentage of reads normalized by the length of the transcript to which they map. The start codon and stop codon regions are most abundant since they are short and contain a large proportion of reads. **(D)** Analysis of footprint distribution for the uORF containing transcript *EIF4G2* only reveals a higher number of reads in the 5’-UTR, compared with the full transcriptome, as expected. **(E)** IGV browser view of the uORF containing *EIF4G2* transcript confirms translation of a uORF, as demonstrated by footprints in the 5’-UTR. Range of y-axes is 10 RPM.

#### Generation of the RNA occupancy files (ROCC)

Following bowtie alignment, a central step in the ribofootPrinter pipeline is to create data structures (ROCC files) to hold the RNA occupancy data from SAM files by using the *builddense* script (Figure 1C). These structures are created using Python dictionaries, with keys for each gene, to include information, such as gene name, sequence, UTR positions, and normalized (rpm) footprint data for a size range of interest (*i.e.* 25-34 nt). In this way, these occupancy files keep metadata and experimental data together. As with other tools (Dunn and Weissman 2016), information can be saved for as 5’ or 3’ end assignment of mapped reads, as well as the coverage of the read (*i.e.* counts across the entire read). While we anticipate most applications will use single-end reads, we also provide an option for mapped paired-end reads (*i.e.* standard mRNA-Seq data). As shown previously, 3’ end assignment often offers superior reading frame alignment for ∼16-nt or ∼28-nt reads from ribosomes in yeast cells, while 5’ end alignment is superior for 40S subunits in yeast or 80S ribosomes in many human cell types (Guydosh and Green 2017; Karasik et al. 2021; Young et al. 2021).

#### Individual transcript analysis

With these ROCC files in hand, several tools are available for analysis (Figure 5A). One of the most basic applications is to output read occupancies as a function of nucleotide position for a particular gene model. This is accomplished with ribofootPrinter’s *writegene2* script. The output data in RPM is provided in csv format which can then be used to create images of ribosome occupancy on a spliced transcript. Like the spliced transcript files created for use in IGV (BEDGRAPH or BIGWIG, described above in Figure 1C) and other transcript-centric viewers (Kiniry et al. 2019), this output offers the advantage over traditional genome viewers in that it excludes intronic regions (Figure 5B, example of *ACTB* gene model). However, ribofootPrinter’s *writegene2* offers the advantage of avoiding artifacts due to averaging performed by IGV’s default windowing function. Comparison between the gene model for *ACTB* produced by *writegene2* (Figure 5B, top track) and the IGV track revealed peak height differences when using IGV’s default *mean windowing function* setting (Figure 5B, center track). If the transcript is too long to display a single peak for each position, this setting causes all peaks within a certain window to be averaged, presumably as a way to improve performance. While the *windowing function* in IGV can be changed to *none* (Figure 5B, bottom track), this setting introduces different artifacts by displaying downsampled data. Overall, ribofootPrinter’s *writegene2* reliably outputs the RPM for each position of the transcript without artifacts that are often introduced by genome browsers.

**Figure 5.**
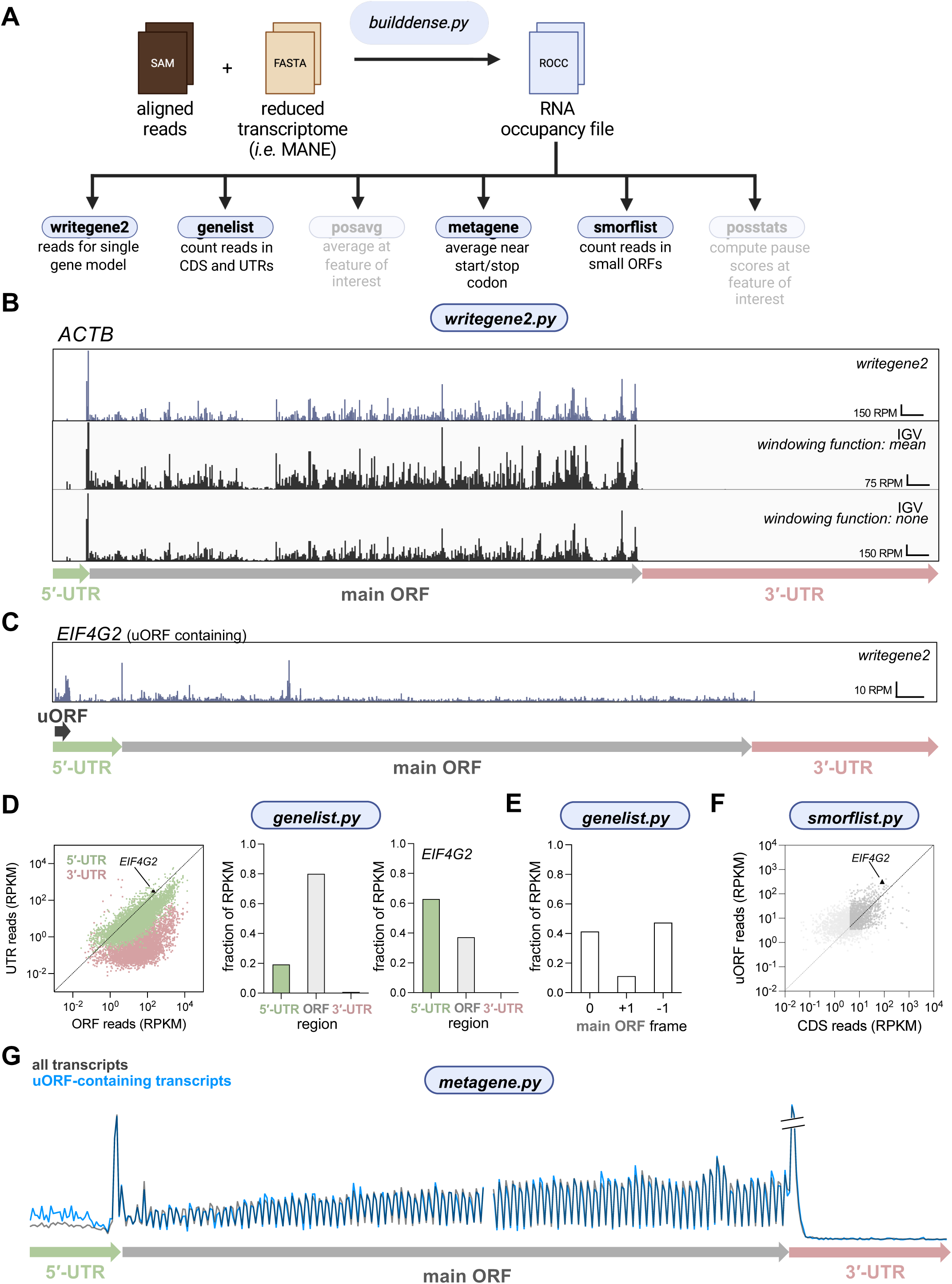
ribofootPrinter can be used to report and analyze reads from across full transcripts. **(A)** Schematic representation of the *writegene2, genelist, metagene* and *smorflist* scripts. These packages use a ROCC file as input. **(B)** Example output of mapped ribosome profiling reads on a gene model for *ACTB* generated by *writegene2* (shown in dark purple). The data are compared against IGV browser views and different windowing settings which influences the representation of the data (shown in black). Range of y-axis is 150 RPM (top), 75 RPM (middle), or 150 RPM (bottom). **(C)** Example output of mapped ribosome profiling reads on the uORF-containing *EIF4G2* transcript generated by *writegene2*. Range of y-axis is 10 RPM. **(D)** Output of *genelist* shows that 5’-UTRs typically have more reads than 3’-UTRs (each dot is a single individual transcript). uORF-containing transcript *EIF4G2* is highlighted as it contains more 5’-UTR reads compared to main ORF reads (*left*). Analysis of mRNA regions reveals the vast majority of reads map to the main ORF with a low-level mapping to the 5’-UTR and even less to the 3’-UTR (*center*). Analysis of the uORF containing *EIF4G2* transcripts reveals a larger number of 5’-UTR reads than all transcripts, as expected and consistent with **C** (*right*). **(E)** The *genelist* script identifies the reading frame of the end of mapped reads. **(F)** Result of small ORF analysis using *smorflist* (each dot is a single predicted uORF) in 5’-UTRs reveals that, as a function of CDS (main ORF) reads, density of reads on individual uORFs is comparable to the CDS (dots more or less along diagonal). This analysis of individual uORFs contrasts with the result in (**D**) where analysis of entire 5’-UTRs was performed. **(G)** Metagene plot for start and stop codons reveals 3-nt periodicity in the coding sequence. It also shows low level translation in the 5’-UTR and nearly no translation in the 3’-UTR, consistent with **D**. Data from a subset of predicted uORF-containing transcripts are overlayed (blue) and show increased translation in the 5’-UTR, as expected. Note that 5’ ends are mapped in **B-G**.

RibofootPrinter’s *genelist* script includes the ability to quantitate total read counts per transcript, reported as both raw reads for differential expression analysis tools, such as DESeq2 (Love et al. 2014), or normalized to total mapped reads and length (in kilobases) of gene models (RPKM) (Figure 5A). Outputs are offered for both coding sequence (ORF) and UTR regions to give a quick portrait of the level of 5’ and 3’-UTR translation levels, values that are known to change due to alterations in the fidelity of initiation or ribosome recycling, respectively. For example, since *EIF4G2* contains a uORF, it is expected to contain a larger number of reads in the 5’-UTR compared to a uORF-less transcript (Figure 5C, Figure 4D, E). Globally, our Ribo-seq sample data reveal that length-normalized footprint densities in 5’-UTRs are considerably greater than in 3’-UTRs as is expected, since under normal conditions more uORF translation occurs compared to stop codon readthrough and consequent 3’-UTR translation (Figure 5D) (Mudge et al. 2022). As expected, *EIF4G2* exhibited a higher proportion of 5’-UTR reads per unit length compared to the entire population (middle and right panels on Figure 5D). In addition, an optional mode outputs reads in each of the 3 reading frames of the coding sequence, allowing visualization of possible frame defects that can result from events such as frameshifting or leaky scanning (Figure 5E). Note that use of this feature assumes the footprint length distribution is narrow enough to distinguish between frames. Factors such as insufficient or inconsistent RNase treatment across samples can limit the utility of this analysis. In general, which frame is considered “0” relates to the “shift” value that estimates the distance between the 5’ or 3’ end of the read and internal features, such as the ribosome A site (see section below on the *metagene* function for further discussion).

#### Small ORF analysis

Accurate quantitation of ribosome footprints on small open reading frames outside coding sequences represents an active area of investigation. These upstream open reading frames (uORFs) in 5’-UTRs and downstream open reading frames (dORFs) in 3’-UTRs are emerging as important regulatory elements that can either enhance or inhibit translation of the main coding sequence (Hinnebusch et al. 2016; Lin et al. 2019; Wu et al. 2020). While metagene analysis (see below) and total quantitation of reads across a UTR region can offer some insight about translation in UTRs, precision estimation of a particular small ORF’s usage is more complicated since small ORFs tend to be overlapping and may initiate with multiple start codons. In addition, initiation may occur with a near-cognate start codon (1-bp mismatch from ATG). In ribofootPrinter’s *smorflist* script (Figure 5A), quantitation of reads on small ORFs is implemented by counting in-frame reads only on all potential small ORFs, thereby excluding most reads from overlapping out-of-frame ORFs. The small ORFs that are analyzed can be filtered by start codon (perfect match to ATG or 1-bp mismatch) and by length, since longer ORFs will be less subject to noise. The output of this analysis can then be combined with appropriate thresholding to identify potentially expressed small ORFs. In a sample dataset, ribosome density across uORFs reveals many uORFs with expression levels comparable to that of the main ORF, including that within *EIF4G2* (Figure 5F).

#### Metagene analysis

The ROCC file offers the ability to perform average (*metagene* function) analysis at start or stop codons of coding sequences (Figure 5A). The script calculates the average read count at a range of nucleotide positions across transcripts that are aligned by their start or stop codon (Supplementary Figure 4A). The average peak that is created at start codons is useful for estimating the position of the ribosome P site with respect to either the 5’ or 3’ end of mapped footprints. This “shift” value is then used by other analysis tools in the package. Typically, 5’ assigned ribosome profiling in either yeast or human cells results in footprint peaks 12 nt upstream of start codons, showing that the first base of the P site is about 12 nt away from the 5’ end (Supplementary Figure 4B).

In addition, metagene analysis for Ribo-seq data also offers a visual metric of reading frame fidelity by revealing how strong the periodicity is in the coding region (Figure 5G). As noted above, this periodicity is also a function of factors that affect the read length distribution, such as RNase treatment. It can also visually reveal the level of translation in UTRs and directly show how it compares with that in the main ORF. Comparison of a metagene computed solely from the previously identified uORF-containing transcripts in Figure 5F (using an available input parameter to filter for specific transcripts) to all transcripts clearly reveals transcripts with uORFs have more ribosomes in their 5’-UTRs (relative to the main ORF) (Figure 5G). In addition, metagene plots computed at stop codons reveal a peak about 10 codons upstream of the stop codon. This peak is likely formed by ribosomes that stall behind the ribosome at the stop codon and are therefore consistent with disomes forming at this position.

Traditional metagene analysis is performed using an arithmetic mean to compute the average (setting: *equalweighting 0*, units of rpm). However, this gives more weight to the most highly expressed genes in the transcriptome and can result in artifacts. Therefore, an alternative with equal weighting of all genes is made available (setting: *equalweighting 1*, units of fraction of reads). Weighting is performed by counting total reads within the coding sequence of the gene (subject to thresholding options to filter out genes with poor read depth) and then normalizing the data to this count prior to averaging. Note that this setting is not recommended for use with more specialized analysis, such as 40S profiling datasets, where the coding sequence cannot be used to evaluate expression level.

#### Metasequence analysis and pause scores (posavg and posstats)

Computing the average read level around sequence-specific positions within gene models (“metasequence” or “metacodon” analysis) is critical for estimating levels of ribosome stalling during elongation. It can also reveal usage of small ORFs in UTRs (uORFs and dORFs) or internal to coding sequences (iORFs). To accomplish this, ribofootPrinter’s *posavg* script accepts a specific nucleotide or amino-acid sequence and then computes the average ribosome footprint level within a defined window around each occurrence of this motif in the transcriptome (Figure 6A). Averages can be subject to thresholding to filter out occurrences with poor read depth. This analysis can be used to check particular motifs for their tendency to slow elongation in coding sequences. For example, when footprints are averaged with a single Pro vs Ser codon are in the P site, Pro clearly slows the ribosome (Figure 6B) more than Ser, as is expected (Schuller et al. 2017).

**Figure 6.**
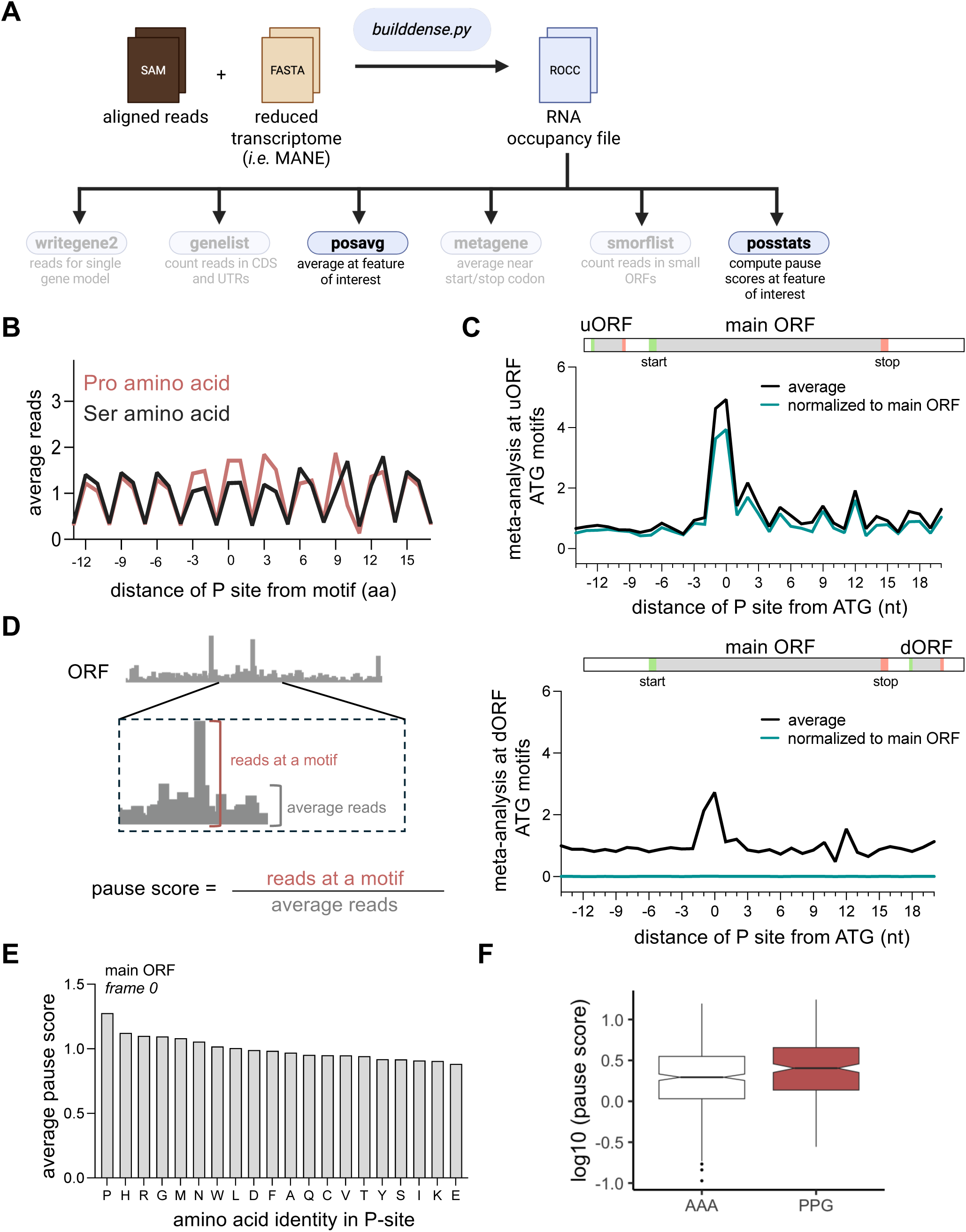
ribofootPrinter can be used to compute and analyze pause scores. **(A)** Schematic representation of the *posavg* and *posstats* scripts. These packages use a ROCC file as input. **(B)** Average analysis using *posavg* of reads at proline (Pro) and serine (Ser) codon motifs reveals a greater peak when a codon encoding a Pro amino acid (but not Ser) is placed in the P site. **(C)** Metaposition analysis of AUG codons in the 5’-UTR (*top*) reveals a clear peak and increased ribosome occupancy and 3-nt periodicity after the peak. This result is consistent with efficient translation of uORFs. Whether data is normalized to a local window (black) or to the respective main ORF sequence (turquoise color) does not dramatically affect the result since uORF translation levels are comparable to main ORF translation levels. Analysis of AUG codons in 3’-UTRs (*bottom*) shows a similar peak when reads are combined with a locally-normalized average, showing dORF translation does take place. However, the lack of a peak when reads are normalized to the main ORF (turquoise color) shows the translation level is quite low. **(D)** Schematic representation of pause score calculations. **(E)** Pause scores from positional average analysis (*all_1* motif setting) at every amino acid reveals Pro induces the strongest stalling. **(F)** Comparable analysis of the distribution of all pause scores at AAA and PPG tri-amino acid motifs shows a difference in the overall distributions with more pausing on PPG.

The *posavg* script allows the average to be locally normalized as a probability density within the window around the motif of interest or normalized to the level of reads in the gene model’s respective coding sequence. For analysis of uORFs (choosing ATG as the motif), the outcome is similar with both methods since ribosome occupancy on uORFs is often comparable to that of coding sequences (Figure 6C, top). In contrast, ribosome occupancy on dORFs is quite low compared with coding sequences and therefore the locally normalized analysis clearly reveals translation on dORFs while the CDS-normalized analysis shows an undetectable level of translation (Figure 6C, bottom). In this way, this metasequence analysis allows a way to check whether there is a change in small ORF translation. In addition, the code can compute a pause score from the meta-analysis plots. The score is computed by dividing the counts in a fixed region around the motif of interest by a local background level defined by inputs to the script (Figure 6D). Analysis for every single amino acid in the ribosome P site clearly shows how Pro is the slowest amino acid on average (Figure 6E), as expected.

Finally, to gain insight about the distribution of pause scores, pause scores at individual motifs of interest can be computed by using the *posstats* script. It computes the pause score by dividing the read count in a narrow window around the position of the motif by the overall level of reads in the region of interest (*i.e.* main ORF or UTR) subject to thresholding of background levels. In general, this script is best run on deep datasets and is most useful for finding particularly large peaks, for example, the strongest examples of ribosome stalling for a specific motif (listed by position within mRNAs). This analysis enables advanced statistical analysis of the distribution of pause scores at motifs of interest and can show global trends. For example, it reveals that the median pause score at Pro-Pro-Gly is higher than Ala-Ala-Ala (Figure 6F) but that the spread in values in considerable.

#### 3D metagene analysis

The ribofootPrinter package can also carry out metagene analysis at start or stop codons as a function of read length. The 3-dimensional (3D) metagene plot that is created by this analysis reveals how different species in a population (*i.e.* ribosomes protecting different footprint sizes) are distributed across a transcript. When performed for both 5’ and 3’ assignment, these plots can reveal how a given ribosome footprint straddles particular features, such as the start codon. Unlike the code that generates a conventional metagene plot, this tool works directly on SAM files, rather than ROCC files, due to the large amount of data required to represent every read in a sample (Figure 7A). Similar to a conventional metagene plot, the location of the 5’ or 3’ end of the mapped read, around start or stop codon, is plotted on the x-axis. The footprint length is then plotted on the y-axis and the read abundance is shown as a heatmap (high abundance in yellow, low abundance in dark purple), thus using 3 dimensions. It also outputs a matching conventional metagene plot. This method can reveal whether the spread in read sizes is primarily due to variation in the position of the 3’ end of the read (Figure 7A *top graphs*) or the 5’ end of the read (Figure 7A *bottom graphs*). In the case of ribosome profiling, this method could therefore distinguish models where proteins that bind to the leading vs. trailing end of a ribosome lead to a longer footprint.

**Figure 7.**
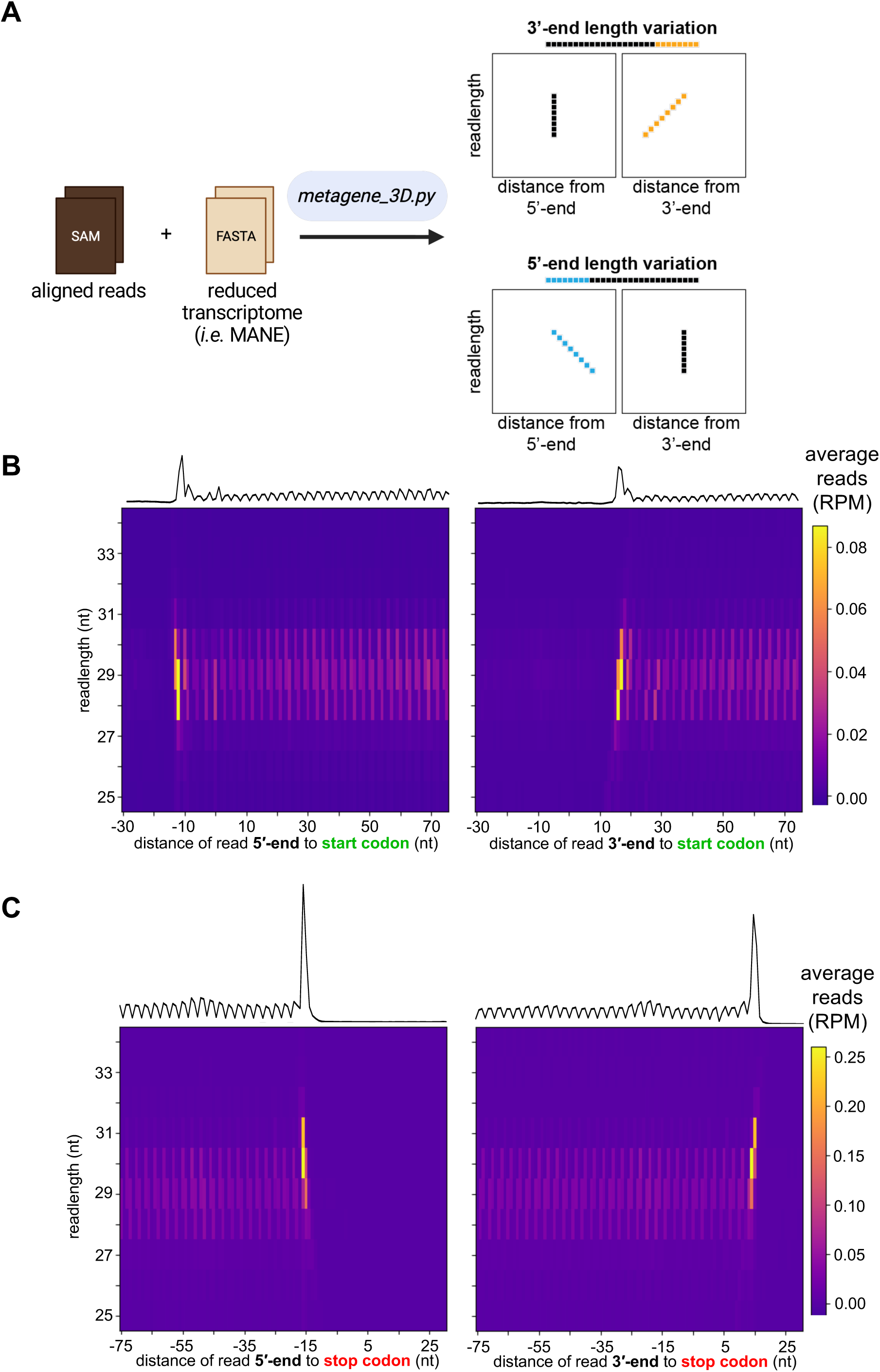
3D metagene analysis provides additional information on read length. **(A)** Schematic representation of the *metagene_3D* script. This script uses a SAM file as input. Graphs explain how 5’-vs 3’-variation is visualized with this analysis. **(B,C)** Comparison of the 3D metagene around start **(B)** or stop **(C)** codon with 5’-(*left*) and 3’-end (*right*) assigned reads. The read abundance in RPM is represented as a heatmap with high abundance colored yellow and low abundance purple (see key on right). Plots demonstrate periodicity and read length variation. For comparison 1-D metagene plots are also generated by this script above 3D plots. Plots are made with *metagene_3D_plot* script.

In the 5’-assigned 3D metagene plot for our Ribo-seq dataset, we observed that the ribosomes that accumulate on start and stop codons in a conventional metagene (Figure 7B, *top* trace) are discernable in the 3D metagene plot as yellow regions (Figure 7B and 7C), consistent with the ribosome footprint abundance analysis (see Figure 4C).

#### Other profiling methods

As noted above, ribofootPrinter is compatible with other types of ribosome profiling preparation methods and other types of short-read sequencing. We therefore examined the performance of the package with 40S subunit profiling (TIS-seq or 40S profiling). First, we determined the footprint length distribution as a probability density (Figure 8A left). We then normalized the length distribution by read count, and read count and feature length (Figure 8A center/right). Unlike the 80S footprints, 40S footprints protect a different distribution of sizes with a peak around 32 nt and long tails. Similar to our 80S dataset, we found that abundance (of reads normalized by read counts) reports a large absolute number of footprints within the ORF (Figure 8A center). Abundance normalized by read counts plus feature length, in contrast, shows how the density of 40S subunits per unit mRNA length is greatest at start and stop codons, followed by the 5’-UTR (Figure 8A, right). This suggests 5’-UTR scanning is relatively quick and that the 40S dwells longer at these codons, perhaps due to slower 60S association and 40S removal.

**Figure 8.**
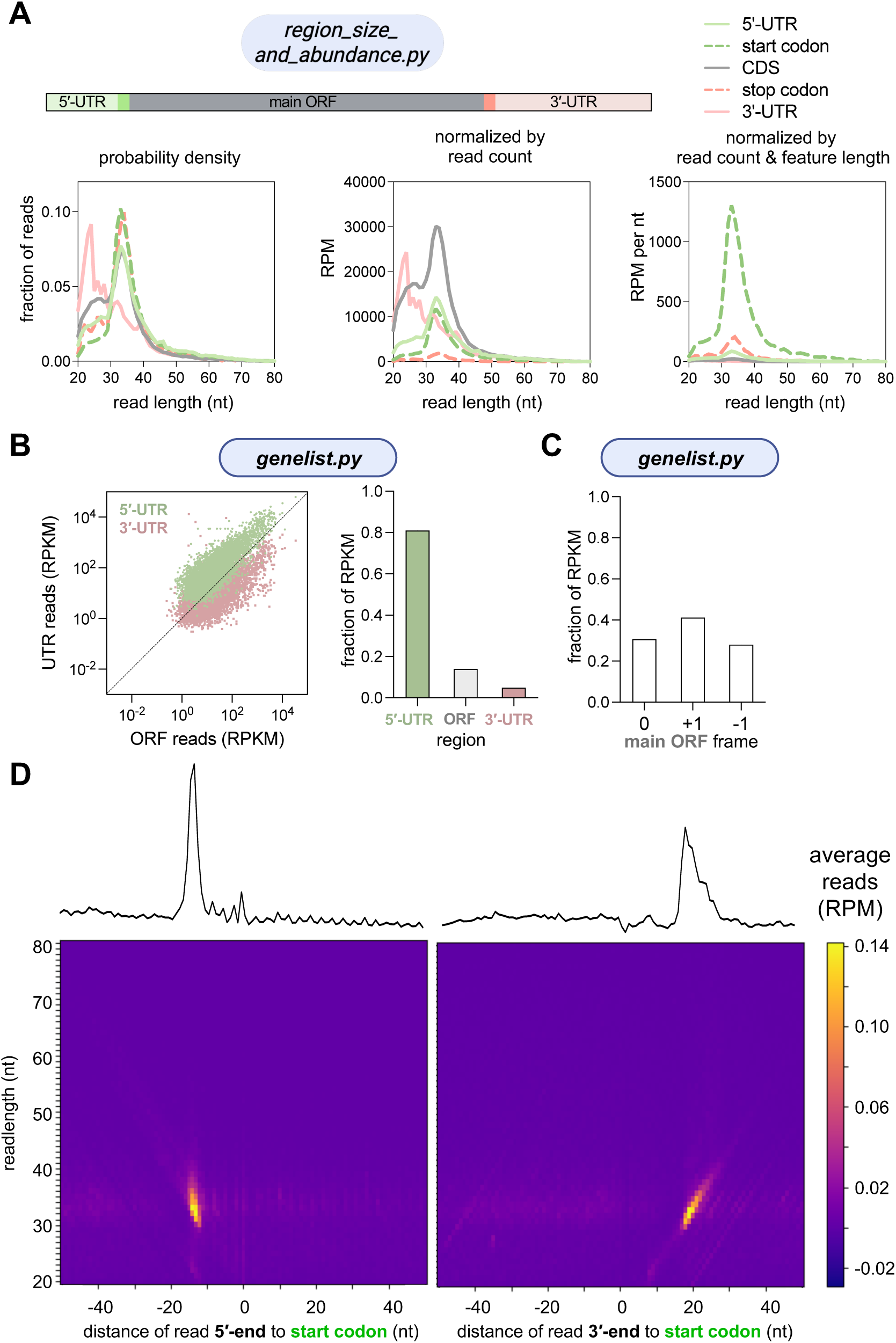
ribofootPrinter is compatible with 40S profiling and other datasets. **(A)** Read length distributions were computed with *region_size_and_abundance* and normalized by using abundance outputs. On the left, the probability density is shown for each region. In the center, the scaling is normalized by overall read abundance in each region. On the right, the histograms are scaled by the abundance and the length of the gene’s feature (*i.e.* 5’-UTR or stop codon) to which each read mapped. **(B)** Output of *genelist* plotted at the level of gene models shows that, as expected, 5’-UTRs of 40S profiling datasets typically have more reads than the main ORF and 3’-UTR (each dot is an individual transcript) (*left*). Analysis of mRNA regions reveals the vast majority of reads map to the 5’-UTR, a lower level map to the main ORF, and even less to the 3’-UTR (*right*). **(C)** The *genelist* script identifies no specific reading frame preference for 40S footprint ends, as expected. **(D)** Analysis using *metagene_3D* of 40S profiling data at start codons. This reveals that 40S subunits are mostly found on start codons. Note the wide footprint length distribution protected by 40S subunits at the start codon is likely caused by the presence of initiation factors. Comparison between 5’- and 3’-assigned plots (*left* vs *right*) shows that the 3’-assiged ends of reads vary with read length since the peak is less vertical (more diagonal) than for 5’-assigned reads on the left.

Next, we investigated our 40S dataset using the *genelist* script. As expected and consistent with our observations above, we found that the majority of the 40S footprints are located within the 5’-UTR (including start codon) (Figure 8B). In addition, the frame distribution of ORF 40S footprints is more random than for 80S footprints (Figure 8C vs 5E). This indicates that the 40S footprints within the ORF likely result from a process that is not dependent on reading frame, such as scanning. If the footprints resulted from translating ribosomes that had fallen apart during sample preparation, the frame distribution would be less equally represented between the three different frames, as shown for 80S data in Figure 5E.

In addition, we plotted 3D metagenes for the 40S profiling dataset at start codons (Figure 8D). Since 40S profiling captures pre-initiation complexes, the majority of the footprints are expected to be localized within the 5’-UTR and start codon. Consistent with this prediction, we find that most of the initiating 40Ss are present on the start codon (likely awaiting recruitment of the 60S subunit), as shown previously (Bohlen et al. 2020). Comparison of the 5’- and 3’-assigned plots shows that the longer footprints are also enriched at the start codon (>40 nt) and tend to be extended on their 3’-end.

#### Considerations for preparation of other transcriptomes

The MANE reduced transcriptome is a carefully curated set of coding transcripts where a single isoform represents the most biologically relevant transcript of each gene (Morales et al. 2022). Currently, mitochondrial genes are not included in the MANE datasets. In addition, curation of coding transcripts was only completed for human. If desired, a custom transcriptome can be generated for other species, mitochondria, viruses, or other needs to be fully compatible with ribofootPrinter. To generate a custom transcriptome, both sequence information and metadata, such as ORF boundaries and transcript length, need to be obtained to generate the FASTA file required for ribofootPrinter (see methods). For example, a fully annotated genome of respiratory syncytial virus (RSV) is available from NCBI (nucleotide: KT992094.1). Both sequences for the full transcripts and annotated CDS can be downloaded and are sufficient to create the FASTA file.

If these transcriptomes will be used in addition to the human transcriptome for example, it is advisable to generate multimapper identifier files for both transcriptomes to identify any similarities between the two transcriptomes that lead to ambiguous mapping.

## Conclusions

We have introduced ribofootPrinter, a readily configurable software toolbox for the analysis of ribosome profiling data. Its approach of reducing mapping complexity by using a reduced transcriptome represents a practical solution to the challenge of aligning reads. This compromise enables the use of and high-resolution tools for analysis of translation, making it a powerful and easy-to-use resource for researchers in the field.

To address the problem of unique mapping inherent in all short-read alignments (Deschamps-Francoeur et al. 2020; Almeida da Paz et al. 2024), we describe multimapping identifier files to view alongside mapped data in a genome bowser to highlight regions of the transcriptome that are prone to multimapping. In addition, ribofootPrinter’s multiple options for meta analysis normalization offer solutions to a problem where needs differ between applications. While no package serves the need of every user, we believe ribofootPrinter offers a useful balance of simplicity and key functionalities that will make it a good choice for many users. In addition, it offers a simple way to get ribosome profiling footprint occupancy and respective mRNA transcript sequences into standard Python list and string data types to help facilitate the development of new forms of analysis.

We also look forward to future enhancements to improve the capabilities of the software. For example, we look forward to adding reduced transcriptomes for other species. It would also be desirable to include capabilities for finding pause peaks and report the underlying sequences as potential pause-inducing motifs. We also anticipate that the growing understanding that translation outside of coding regions is important for regulating gene expression and generating functional peptides will necessitate additional tools for characterization of ORFs in noncoding regions (Chen et al. 2020). Finally, we expect additional tools could be created for analysis of other types of small RNAs besides ribosome footprints, such as those generated by clip-based approaches. We believe the straightforward design of ribofootPrinter will enable continued growth and adaptation to study these phenomena.

## Methods

ribofootPrinter is available on Github: https://github.com/guydoshlab/ribofootPrinter2. The current release is 2.1, which is also available on Zenodo with DOI: 10.5281/zenodo.17956096

### ribofootPrinter compatibility and packages

The software is best run inside a Python 3 virtual environment (venv) where Biopython (and other required packages, which include matplotlib, pandas, openpyxl) have been installed. Packages required in the environment to create figures in this manuscript are given on Github. We assume the user has appropriately pre-processed ribosome footprint reads, including steps to trim linkers, remove duplicates using unique molecular identifiers (UMIs), and subtract rRNA and other non-coding RNA sequences (Step 1 in Figure 1B). At this stage, the reads can be aligned to the reduced transcriptome of choice using bowtie. In this study, the reduced MANE v1.4 transcriptome (shortnames) FASTA file was used and can be downloaded from Github or created using the instructions provided therein. A set of specialized tools is also required to run the code and is provided in the repository as tools.py.

Once SAM files are obtained from the bowtie alignment, they can be converted into the sequence and read data structures (ROCC files) using the *builddense* function (described below). ROCC files contain important mapping information and serve as input files for *writegene2, metagene, genelist, smorflist, posavg* and *posstats* packages. The packages *region_size_and_abundance* and *metagene_3D* are run directly on the SAM file.

The longnames.fasta file used for running ribofootPrinter can be downloaded from the Github page (https://github.com/guydoshlab/ribofootPrinter2.0-beta/tree/main/preparation/MANE_v1.4_Preparation/output_files) or created using the instructions provided therein. The longnames.fasta file must adhere to the following header format with all fields separated by “|” characters, as shown in this example:

~~~
>ENSG00000111640.15|ENST00000229239.10|ENSP00000229239.5|NM_0020
46.7|NP_002037.2|GAPDH|1285|UTR5:1-76|CDS:77-1084|UTR3:1085-1285
~~~

The first field should be the formal gene accession number, i.e. “ENSG00000111640.15” followed by the (common) gene name in the sixth field (GAPDH in the example). This is followed by overall gene length. The final 3 fields should include the prefixes “UTR5:”, “CDS:”, and “UTR3:” followed by the start and end position of that region, separated by a hyphen.

All analysis routines are performed on the command line by passing relevant parameters to the Python script. These parameters are read by the code and a metadata output, including error messages, and the read-in parameters are printed to screen within the terminal and can be captured for record-keeping requirements. All data outputs are in csv format or matplotlib window for 3D metagenes.

A detailed description of each tool is provided on Github and further details about the input arguments is available in Supplementary Table 1. Similar information is also found in the comments in the code. An executable file of shell commands is provided as a Supplementary File (ribofootprinter.sh) to generate data used for figures in this manuscript.

### Dataset library preparation

We used two different types of datasets to provide examples for ribofootPrinter’s scripts: 80S and 40S ribosome profiling. Libraries for both datasets were generated from A549 cells following a previously described protocol (McGlincy and Ingolia 2017). Prior to library preparation of the 40S samples, the following steps were completed as described previously (Bohlen et al. 2020; Wagner et al. 2022). The cells were crosslinked prior to harvest. Following lysis, the ribosomes were separated on a 15%- 35% sucrose gradient and fractions corresponding to the 40S subunit were collected. The RNA was reverse crosslinked and extracted by acid-phenol-chloroform extraction. Once RNA was obtained, the libraries were prepared similarly according to the standard protocol for Ribo-Seq. Gel-based read size selection for the 40S dataset was 20-80 nt while that for the 80S dataset was 25-34 nt range.

### Dataset library alignment

Once libraries were prepared, they were sequenced as single-end reads on a Novaseq X 1.5B 100 cycle chip. The raw FASTQ files were processed as described (McGlincy and Ingolia 2017) to complete trimming, deduplication, and removal of contaminating non-coding reads (Step 1 in Figure 1B). The footprint-containing FASTQ files were then aligned against the reduced MANE v1.4 transcriptome (shortnames.FASTA) using the following bowtie1 settings (*-v 1 -y -S*), resulting in a SAM file. To reduce the size of the SAM files, each was downsampled by randomly selecting 50% of the reads with the code below.

Example SAM files and ROCC files used to create figures in this manuscript are available on Zenodo (10.5281/zenodo.17917807). In addition, FASTQ files were generated from these downsampled SAM files and are available on NIH SRA (SRR35611919 and SRR35611920).

### Visualizing datasets in IGV

As noted in the main text, a genome browser can be utilized to view the transcriptome aligned reads. We provide a guide on how to convert SAM files into IGV compatible files on the Github page (https://github.com/guydoshlab/ribofootPrinter2.0-beta/tree/main/preparation/MANE_v1.4_IGV). This includes conversion of SAM files into IGV-compatible BAM or BEDGRAPH/BIGWIG files. The GTF file, which outlines the ORF boundaries for MANE v1.4 transcriptome transcripts, is available for download from Github (https://github.com/guydoshlab/ribofootPrinter2.0-beta/tree/main/preparation/MANE_v1.4_IGV/output_files) or can be created using the instructions provided therein. Example BEDGRAPH files are provided on Github.

### Multimapping

Generation of the transcriptome-derived FASTQ files and multimapper identifier files (*mm_id*) used for Figures 2-3 and Supplementary Figure 2 is described in the supplemental computational analysis file (multimapper_code.pdf) using custom Python code and the seqkit package (Shen et al. 2016). The *mm_id_*files are available on Github.

### Clustal Omega protein alignment

Nucleotide sequences in fasta format for *ACTB, POTEI* and *POTEJ* were obtained from the MANEv1.4_longnames.FASTA file. These sequences were aligned using the Multiple Sequence Alignment (MSA) tool from Clustal Omega (Madeira et al. 2024) using settings (Sequence type: *RNA,* Parameters: *ClustalW with character counts*).

## Supporting information

Multimapper code

shell commands

## Acknowledgements

We thank Agnes Karasik for help identifying bugs in early versions of the code and the Guydosh Lab and James Marks for helpful discussions about the vision, development, presentation, and naming of the code. We thank Hernan Lorenzi for feedback on the manuscript. We also thank Shashikant Pujar for helpful discussion about the MANE project. Sequencing was provided by the NHLBI DNA Sequencing Core.

## Supplementary Figure Legends

**Supplementary Figure 1.**
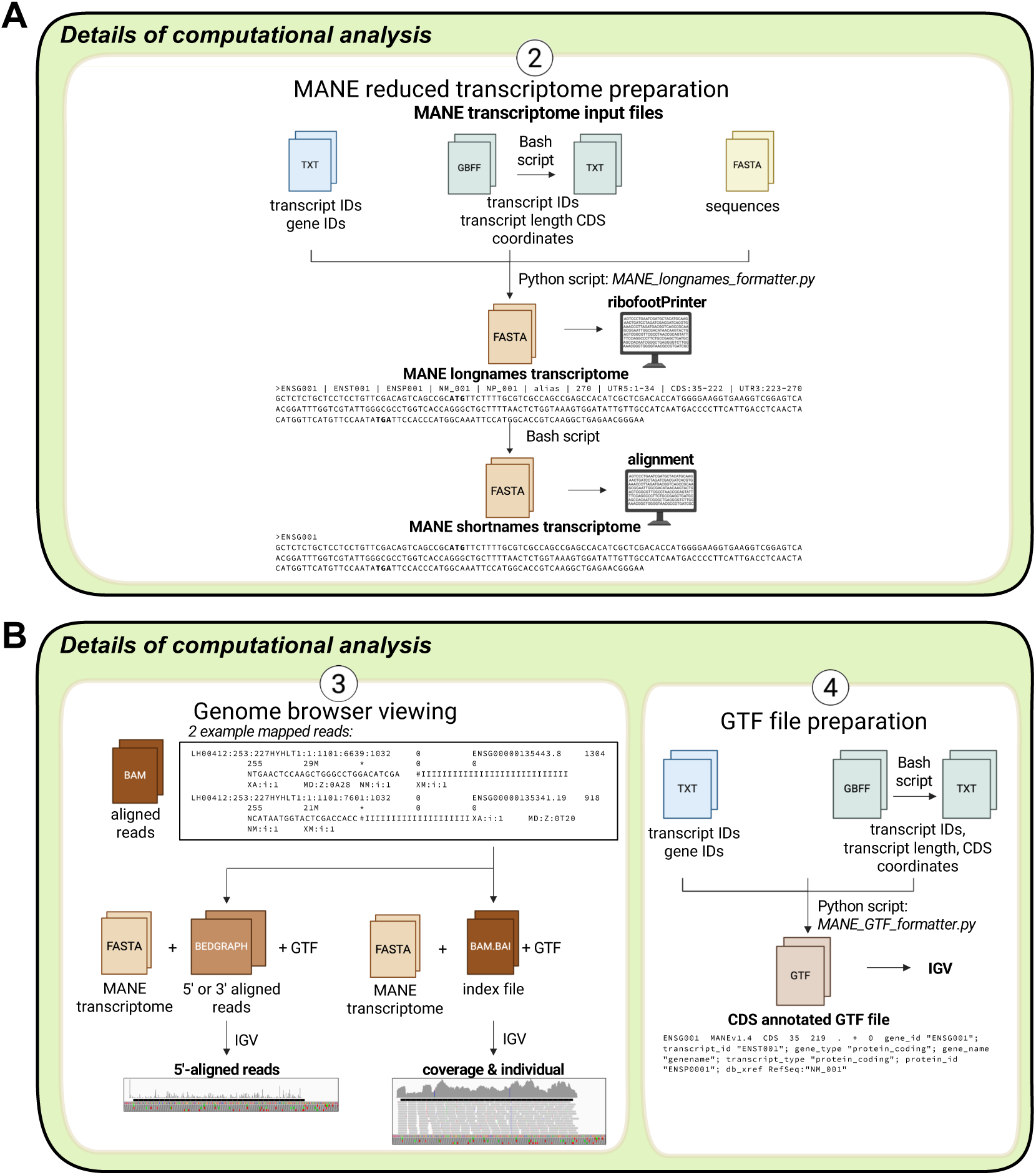
Overview of major steps in read processing prior to running key scripts in ribofootPrinter. **(A)** The MANE transcriptome (FASTA files) for bowtie alignments (shortnames) or ribofootPrinter (longnames) can be created using commands that are provided on Github. **(B)** Description of the pipeline used to convert aligned SAM files into genome browser compatible files (*left*). GTF files for IGV can be downloaded or prepared using the script available on Github (*right*).

**Supplementary Figure 2.**
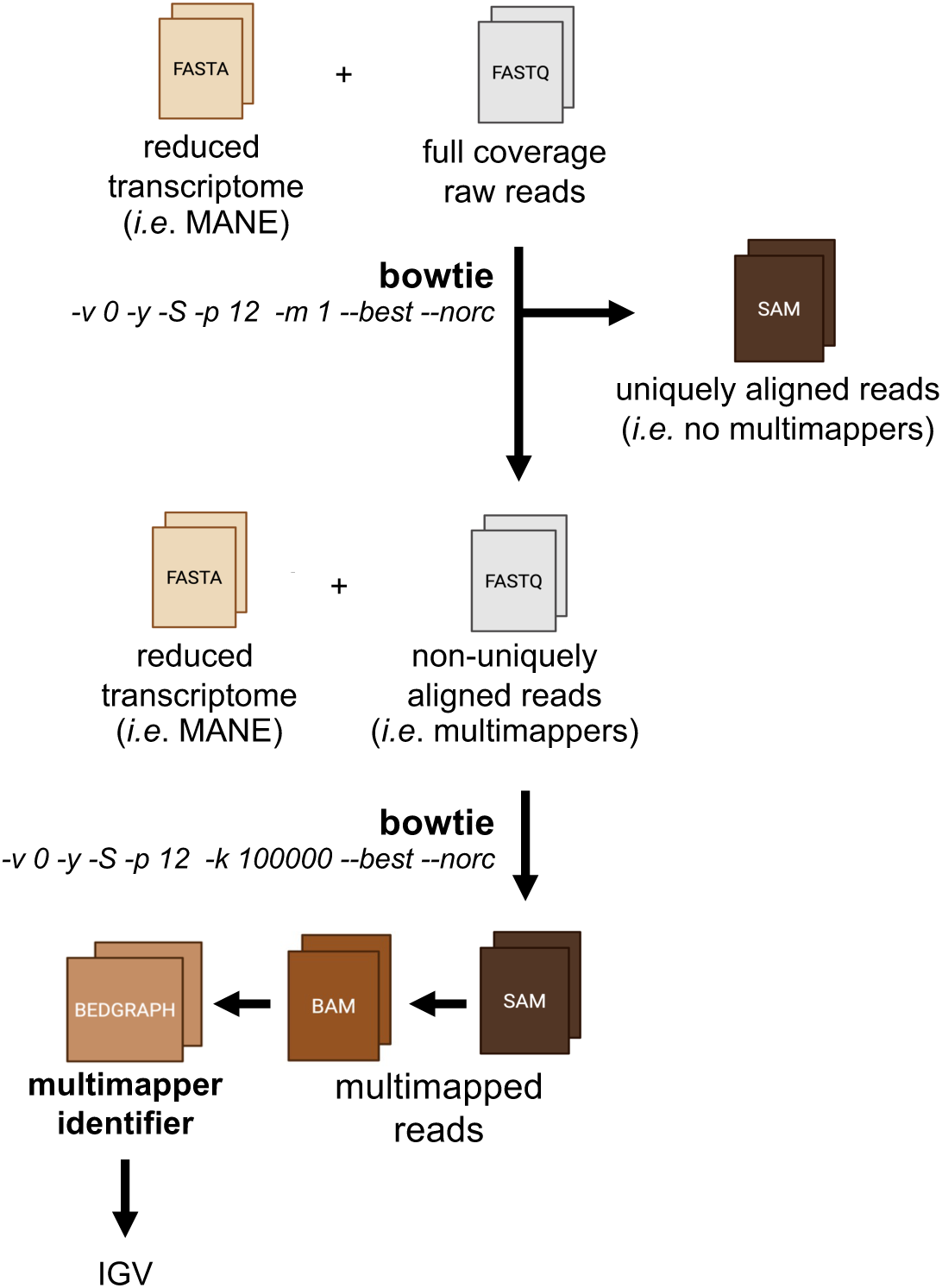
Generation of multimapper identifier (mm_id) files. The transcriptome-derived FASTQ files are aligned against the transcriptome using bowtie settings that only allow uniquely mapped reads (*-m 1*). This results in a SAM file containing uniquely mapped reads only and a FASTQ file with unmapped reads (containing multimappers). The FASTQ file containing non-uniquely aligned reads is aligned against the reduced transcriptome again with bowtie using a setting allowing a large number of multimapping events (*-k 100000*). This generates a SAM file containing multimapped reads which can be converted into IGV compatible BEDGRAPH files termed multimapper identifier files.

**Supplementary Figure 3.**
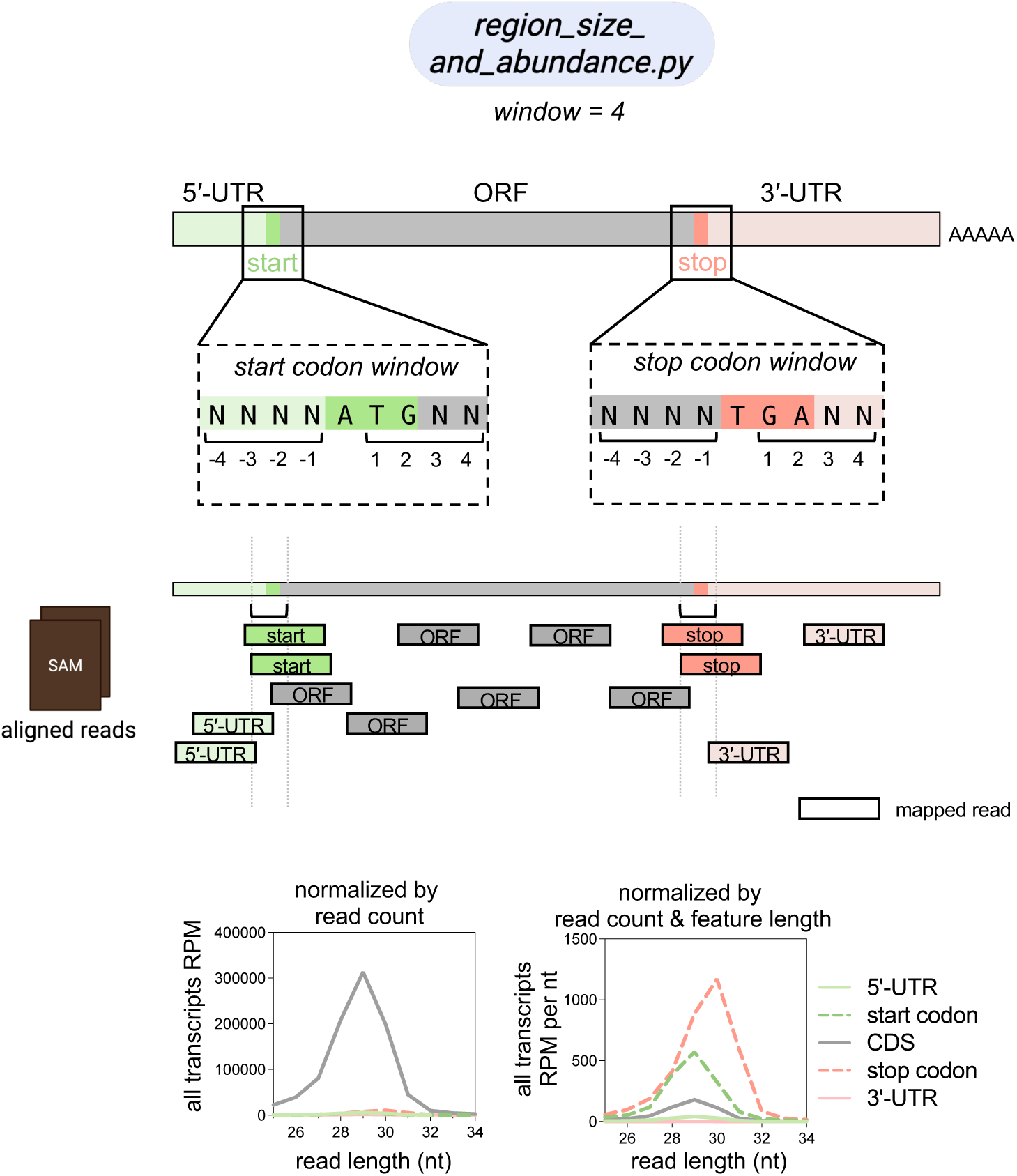
Algorithm used to determine footprint length distribution within UTRs, main ORF, start codon or stop codon. The *region_size_and_abundance* script defines a window around the start and stop codons which will determine the UTR and main ORF boundaries. A read is counted as start or stop if the read overlaps with the start codon or stop codon region. Only partially overlapping reads are distributed into the UTR or main ORF, depending on their location. This is shown as a cartoon (*top*). The two abundance calculations output by the script can be used to normalize the output probability density histogram and create histograms with alternative normalization (*bottom*).

**Supplementary Figure 4.**
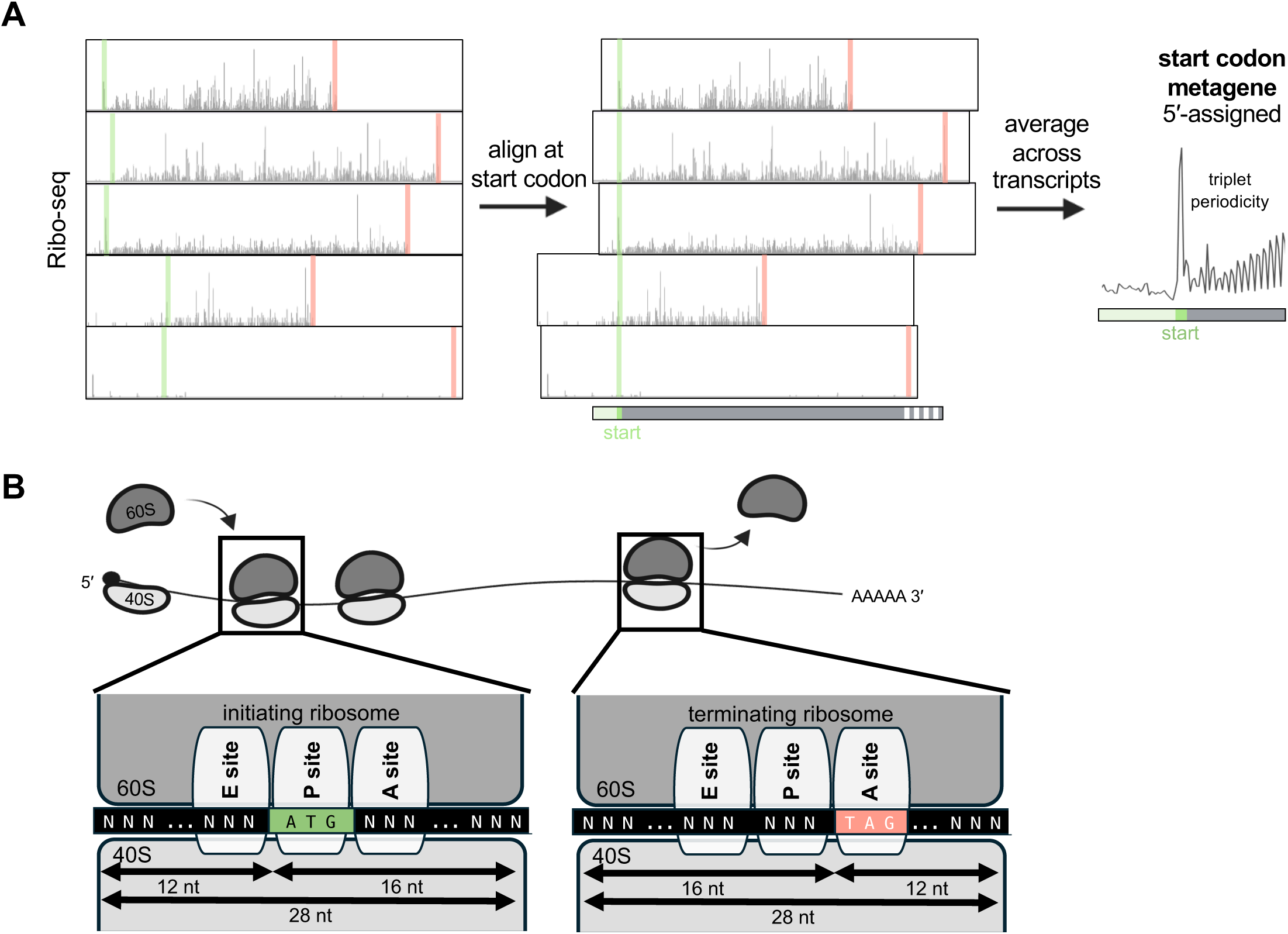
Algorithm used to generate metagene plots. **(A)** Metagene plots are generated by aligning the mapped reads around the start or stop codon, followed by taking the average across all transcripts. This reveals nucleotide periodicity within the main ORF for Ribo-seq data. **(B)** Schematic representation of the ribosome protecting a fixed number of nucleotides (this can differ depending on the sample preparation method). Different types of analysis can be completed for 5’- or 3’-end assigned reads. If desired, shifts can be calculated to study the E-, P- or A- sites.

