## Supplementary material for "ribofootPrinter: A precision python toolbox for analysis of ribosome profiling data": Multimapper code

***Generation of transcriptome-derived FASTQ files and analysis with bowtie***

Transcriptome-derived FASTQ files containing every possible read that could be derived from the transcriptome sequences were generated using a custom Python script (readgenerator_fullcov.py) that is available on Github (https://github.com/guydoshlab/ribofootPrinter2). This script used the MANEv1.4_longnames reduced transcriptome file as input file and outputs FASTQ files with different read lengths as the setting *readlength* is adjusted. We checked that the reads in the FASTQ files fully covered the transcriptome by visualizing them (as BAM files generated by bowtie) in IGV (for example, Figure 2C). The number of unique reads within the full coverage transcriptome-derived FASTQ files (Figure 2C) was determined using the seqkit package (Shen et al. 2016).

Next, the full coverage transcriptome-derived FASTQ files (named *.fq) were aligned against the shortnames transcriptome using different settings to determine the effect of read length on multimapping events. It is assumed that the bowtie executable file is in the local directory or is in the path. Note the output statistics are used for analysis for this code; output files can also be viewed in IGV if converted to BAM format.

```
mkdir -p nomatch
REF=/filepath/bowtie_indexed/MANE_transcriptome/MANEv1.4
for file in *.fq; do echo $file; bowtie -v 0 -y -S -p 12 -k 2 --
best --un ./nomatch/${file%.fq}_nomatch.fq -x $REF ./$file
./${file%.fq}.sam; done
```

The number of mismatches can be changed by adjusting *-v*. The -S setting outputs a SAM file. It is important to include the -k 2 setting which allows bowtie to report up to 2 alignments which is needed to identify multimapping events. Following alignment, bowtie will output information about read alignment. Subtracting the reads with at least one alignment from “reported alignments” gives the number of multimapped reads (this is possible because we only allow mapping to one other site by using the -k 2 setting). The percentage multimapped reads (y-axis in Figure 2D) can be calculated by dividing the number of multimapped reads by reads processed. It is not expected that any files end up in the nomatch folder with this -k 2 setting.

***Generation of multimapper identifier files (mm_id)***

The full coverage transcriptome-derived FASTQ files (named *.fq) were used to identify multimapping regions. It is assumed that the bowtie executable file is in the local directory or is in the path. Note that the analysis below will overwrite files from the analysis in the previous section if the environment the same. First, the FASTQ files were aligned against the reduced transcriptome using bowtie:

```
mkdir -p nomatch
REF=/filepath/bowtie_indexed/MANE_transcriptome/MANEv1.4
for file in *.fq; do echo $file; bowtie -v 0 -y -S -p 12 -m 1 --
best --norc --un ./nomatch/${file%.fq}_nomatch.fq -x $REF
./$file ./${file%.fq}.sam; done
```

The parameter *-m 1* instructs bowtie to only report unique alignments which allows us to divide the reads between uniquely mapped reads (exported as a SAM file, -*S* setting) and multimapped reads (exported as a nomatch FASTQ file, --*un ./nomatch/${file%.fq}_nomatch.fastq*). Once the nomatch FASTQ file containing multimapped reads has been obtained, they are realigned to the reduced transcriptome with bowtie settings allowing multiple multimapped reads *(-k 100000* setting). This will output a SAM file containing information on exclusively multimapped reads.

```
cd nomatch

REF=/filepath/bowtie_indexed/MANE_transcriptome/MANE_v1.4
for file in *.fq; do echo $file; bowtie -v 0 -y -S -p 12 -k
100000 --best --norc -x $REF ./$file ./${file%.fq}.sam; done
```

The SAM file containing multimapped reads is converted into 5'-end aligned bedgraph files using the *samtools* and *bedtools* package.

```
for file in *.sam; do samtools sort -o ${file%.sam}.bam $file;
done

for file in *.bam; do echo $file; bedtools genomecov -ibam $file
-bg -5 >  ${file%.bam}_mm_id.bedgraph; done
```

These multimapper identified (mm_id.bedgraph) files can be viewed in IGV together with aligned reads of interest. The metric encoded in these files is therefore 0 for positions that map uniquely. Otherwise, the value indicates the number of sites the read could map to, and is capped at, 100,000 sites. Since most cases of multiple mapping involve a few sites, it is advisable to set the axis limits to <10 when viewing these files.
