## Supplementary material for "ribofootPrinter: A precision python toolbox for analysis of ribosome profiling data": shell commands: ribofootprinter.sh

```
#!/bin/sh

# Guidance on use of this file:
# Download ribofootPrinter from Github.
# Following ribofooPrinter download, navigate to the folder containing the code. 
# For example: cd ./ribofootPrinter2-2.1.0
# Make sure this .sh file is in this folder.
# Then create a folder called sam_files. In it, place the zipped sam files.
# Zipped samfiles are available here: 
# https://zenodo.org/records/17917807/files/40S.SAM.zip
# https://zenodo.org/records/17917807/files/80S.SAM.zip
# Make sure you use a virtual environment with needed dependencies (see Github). 
# Run this .sh file. Note you may need to make it executable using the chmod command.

ls

# Running the *ribofootPrinter* package
## Generate folders for data files
mkdir -p ./rocc_files/
mkdir -p ./data_files/ 

## Create variables for different paths
FILEPATH=$(pwd)

CODE=/$FILEPATH/code
FASTA=/$FILEPATH/preparation/MANE_v1.4_Preparation/output_files
SAM=/$FILEPATH/sam_files
ROCC=/$FILEPATH/rocc_files
DATA=/$FILEPATH/data_files
FILES=/$FILEPATH/associated_files

mkdir -p ./data_files/writegene2 ./data_files/metagene ./data_files/genelist ./data_files/smorflist ./data_files/posavg ./data_files/posstats ./data_files/region_size_and_abundance ./data_files/metagene_3D


## Unzip files
for file in $FASTA/*.zip; do unzip $file; done
for file in $FILES/*.zip; do unzip $file; done
for file in $SAM/*.zip; do unzip $file; done

mv *.fasta $FASTA
mv *.ebwt $FASTA  
mv *.xlsx $FILES 
mv *.SAM $SAM

echo "Directories have been set up and files unzipped."


## SCRIPT 1. Run *builddense* to generate ROCC files from SAM files
python $CODE/builddense.py $FASTA/MANEv1.4_longnames.fasta $SAM/80S.SAM $ROCC/80S -1 25 34 1 > $ROCC/80S_bd_metadata.txt 
python $CODE/builddense.py $FASTA/MANEv1.4_longnames.fasta $SAM/40S.SAM $ROCC/40S -1 20 80 1 > $ROCC/40S_bd_metadata.txt 

if [ ! -f $ROCC/80S.rocc ]; then echo "ERROR with running builddense on 80S dataset. Files not created."; else echo "builddense finished running. 80S.ROCC files have been created."; fi

if [ ! -f $ROCC/40S.rocc ]; then echo "ERROR with running builddense on 40S dataset. Files not created."; else echo "builddense finished running. 40S.ROCC files have been created."; fi


## SCRIPT 2. Run *writegene2* from ROCC files
python $CODE/writegene2.py $ROCC/80S.rocc ACTB $DATA/writegene2/80S_wg2_ACTB > $DATA/writegene2/80S_wg2_ACTB_metadata.txt
python $CODE/writegene2.py $ROCC/80S.rocc EIF4G2 $DATA/writegene2/80S_wg2_EIF4G2 > $DATA/writegene2/80S_wg2_EIF4G2_metadata.txt

if [ ! -f $DATA/writegene2/80S_wg2_ACTB.csv ]; then echo "ERROR with running writegene2 for ACTB. Files not created."; else echo "writegene2 finished running for ACTB. Data can be found in the data_files folder."; fi

if [ ! -f $DATA/writegene2/80S_wg2_EIF4G2.csv ]; then echo "ERROR with running writegene2 for EIF4G2. Files not created."; else echo "writegene2 finished running for EIF4G2. Data can be found in the data_files folder."; fi


## SCRIPT 3. Run *genelist* from ROCC files
python $CODE/genelist.py $ROCC/80S.rocc 12 1 $DATA/genelist/80S_gl > $DATA/genelist/80S_gl_metadata.txt 
python $CODE/genelist.py $ROCC/40S.rocc 12 1 $DATA/genelist/40S_gl > $DATA/genelist/40S_gl_metadata.txt 

if [ ! -f $DATA/genelist/80S_gl.csv ]; then echo "ERROR with running genelist. 80S files not created."; else echo "genelist finished running for 80S dataset. Data can be found in the data_files folder."; fi

if [ ! -f $DATA/genelist/40S_gl.csv ]; then echo "ERROR with running genelist. 40S files not created."; else echo "genelist finished running for 40S dataset. Data can be found in the data_files folder."; fi


## SCRIPT 4. Run *posavg* from ROCC files
python $CODE/posavg.py $ROCC/80S.rocc all_1 1 0 0 30 0 12 1 none $DATA/posavg/80S_pa_all_1_aa_CDS_frame0 > $DATA/posavg/80S_pa_all_1_aa_CDS_frame0_metadata.txt

if [ ! -f $DATA/posavg/80S_pa_all_1_aa_CDS_frame0_avgdata.csv ]; then echo "ERROR with running posavg all_1 amino acids. Files not created."; else echo "posavg finished running for all_1 amino acids. Data can be found in the data_files folder."; fi

python $CODE/posavg.py $ROCC/80S.rocc ATG 0 3 0 30 0 12 0 none $DATA/posavg/80S_pa_ATG_UTR5_allframe_norm0 > $DATA/posavg/80S_pa_uORF_norm0_metadata.txt 
python $CODE/posavg.py $ROCC/80S.rocc ATG 0 3 0 30 1 12 0 none $DATA/posavg/80S_pa_ATG_UTR5_allframe_norm1 > $DATA/posavg/80S_pa_uORF_norm1_metadata.txt 

if [ ! -f $DATA/posavg/80S_pa_ATG_UTR5_allframe_norm0_avgdata.csv ]; then echo "ERROR with running posavg uORFs. Files not created."; else echo "posavg uORFs finished running. Data can be found in the data_files folder."; fi

if [ ! -f $DATA/posavg/80S_pa_ATG_UTR5_allframe_norm1_avgdata.csv ]; then echo "ERROR with running posavg uORFs. Files not created."; else echo "posavg uORFs finished running. Data can be found in the data_files folder."; fi

python $CODE/posavg.py $ROCC/80S.rocc ATG 0 3 0 30 0 12 2 none $DATA/posavg/80S_pa_ATG_UTR3_allframe_norm0 > $DATA/posavg/80S_pa_dORF_norm0_metadata.txt 
python $CODE/posavg.py $ROCC/80S.rocc ATG 0 3 0 30 1 12 2 none $DATA/posavg/80S_pa_ATG_UTR3_allframe_norm1 > $DATA/posavg/80S_pa_dORF_norm1_metadata.txt 

if [ ! -f $DATA/posavg/80S_pa_ATG_UTR3_allframe_norm0_avgdata.csv ]; then echo "ERROR with running posavg dORFs. Files not created."; else echo "posavg dORFs finished running. Data can be found in the data_files folder."; fi

if [ ! -f $DATA/posavg/80S_pa_ATG_UTR3_allframe_norm1_avgdata.csv ]; then echo "ERROR with running posavg dORFs. Files not created."; else echo "posavg dORFs finished running. Data can be found in the data_files folder."; fi


## SCRIPT 5. Run *metagene* from ROCC files
python $CODE/metagene.py $ROCC/80S.rocc 1 1 5 50 300 none $DATA/metagene/80S_mg_start > $DATA/metagene/80S_mg_start_metadata.txt
python $CODE/metagene.py $ROCC/80S.rocc 2 1 5 300 50 none $DATA/metagene/80S_mg_stop > $DATA/metagene/80S_mg_stop_metadata.txt

if [ ! -f $DATA/metagene/80S_mg_start.csv ]; then echo "ERROR with running metagene. Files not created."; else echo "metagene start finished running. Data can be found in the data_files folder."; fi

if [ ! -f $DATA/metagene/80S_mg_stop.csv ]; then echo "ERROR with running metagene. Files not created."; else echo "metagene stop finished running. Data can be found in the data_files folder."; fi

python $CODE/metagene.py $ROCC/80S.rocc 1 1 5 50 300 $FILES/subset_list.xlsx $DATA/metagene/80S_mg_start_subset > $DATA/metagene/80S_mg_start_subset_metadata.txt
python $CODE/metagene.py $ROCC/80S.rocc 2 1 5 300 50 $FILES/subset_list.xlsx $DATA/metagene/80S_mg_stop_subset > $DATA/metagene/80S_mg_stop_subset_metadata.txt

if [ ! -f $DATA/metagene/80S_mg_start_subset.csv ]; then echo "ERROR with running metagene start subset dataset. Files not created."; else echo "metagene start finished running. Data can be found in the data_files folder."; fi

if [ ! -f $DATA/metagene/80S_mg_start_subset.csv ]; then echo "ERROR with running metagene stop subset dataset. Files not created."; else echo "metagene stop finished running. Data can be found in the data_files folder."; fi


## SCRIPT 6. Run *smorflist* from ROCC files
python $CODE/smorflist.py $ROCC/80S.rocc 4 12 0 0 5 $DATA/smorflist/80S_uorflist > $DATA/smorflist/80S_uorflist_metadata.txt

if [ ! -f $DATA/smorflist/80S_uorflist.csv ]; then echo "ERROR with running smorflist. Files not created."; else echo "smorflist finished running. Data can be found in the data_files folder."; fi


## SCRIPT 7. Run *posstats* from ROCC files
python $CODE/posstats.py $ROCC/80S.rocc PPG 1 0 10 1 12 1 $DATA/posstats/80S_ps_PPG >  $DATA/posstats/80S_ps_PPG_metadata.txt
python $CODE/posstats.py $ROCC/80S.rocc AAA 1 0 10 1 12 1 $DATA/posstats/80S_ps_AAA >  $DATA/posstats/80S_ps_AAA_metadata.txt

if [ ! -f $DATA/posstats/80S_ps_PPG_80S.csv ]; then echo "ERROR with running posstats. PPG files not created."; else echo "posstats finished running. Data can be found in the data_files folder."; fi
if [ ! -f $DATA/posstats/80S_ps_AAA_80S.csv ]; then echo "ERROR with running posstats. AAA files not created."; else echo "posstats finished running. Data can be found in the data_files folder."; fi


## SCRIPT 8. Run *region_size_and_abundance* from SAM files
python $CODE/region_size_and_abundance.py $FASTA/MANEv1.4_longnames.fasta $SAM/80S.SAM $DATA/region_size_and_abundance/80S_rsa 25 34 4 none > $DATA/region_size_and_abundance/80S_rsa_metadata.txt

python $CODE/region_size_and_abundance.py $FASTA/MANEv1.4_longnames.fasta $SAM/80S.SAM $DATA/region_size_and_abundance/80S_rsa_subset 25 34 4 $FILES/subset_list_EIF4G2.xlsx > $DATA/region_size_and_abundance/80S_rsa_subset_metadata.txt

python $CODE/region_size_and_abundance.py $FASTA/MANEv1.4_longnames.fasta $SAM/40S.SAM $DATA/region_size_and_abundance/40S_rsa 20 80 4 none > $DATA/region_size_and_abundance/40S_rsa_metadata.txt

if [ ! -f $DATA/region_size_and_abundance/80S_rsa.csv ]; then echo "ERROR with running region_size_and_abundance 80S dataset. Files not created."; else echo "region_size_and_abundance finished running. Data can be found in the data_files folder."; fi
if [ ! -f $DATA/region_size_and_abundance/80S_rsa_subset.csv ]; then echo "ERROR with running region_size_and_abundance 80S subset dataset. Files not created."; else echo "region_size_and_abundance finished running. Data can be found in the data_files folder."; fi
if [ ! -f $DATA/region_size_and_abundance/40S_rsa.csv ]; then echo "ERROR with running region_size_and_abundance 40S dataset. Files not created."; else echo "region_size_and_abundance finished running. Data can be found in the data_files folder."; fi

## SCRIPT 9. Run *metagene_3D* from SAM files
python $CODE/metagene_3D.py $FASTA/MANEv1.4_longnames.fasta $SAM/80S.SAM $DATA/metagene_3D/80S_3dmg_start none 25 34 30 75 1 > $DATA/metagene_3D/80S_3dmg_start_metadata.txt
python $CODE/metagene_3D.py $FASTA/MANEv1.4_longnames.fasta $SAM/80S.SAM $DATA/metagene_3D/80S_3dmg_stop none 25 34 75 30 2 > $DATA/metagene_3D/80S_3dmg_stop_metadata.txt

python $CODE/metagene_3D.py $FASTA/MANEv1.4_longnames.fasta $SAM/40S.SAM $DATA/metagene_3D/40S_3dmg_start none 20 80 50 50 1 > $DATA/metagene_3D/40S_3dmg_start_metadata.txt

if [ ! -f $DATA/metagene_3D/80S_3dmg_start_1Dmetas.csv ]; then echo "ERROR with running metagene_3D start 80S dataset. Files not created."; else echo "80S metagene_3D start finished running. Data can be found in the data_files folder."; fi

if [ ! -f $DATA/metagene_3D/80S_3dmg_stop_1Dmetas.csv ]; then echo "ERROR with running metagene_3D stop 80S dataset. Files not created."; else echo "80S metagene_3D stop finished running. Data can be found in the data_files folder."; fi

if [ ! -f $DATA/metagene_3D/40S_3dmg_start_1Dmetas.csv ]; then echo "ERROR with running metagene_3D start 40S dataset. Files not created."; else echo "40S metagene_3D start finished running. Data can be found in the data_files folder."; fi


echo "Analysis complete. See terminal output above for overview. "
```
